# Nucleation and spreading rejuvenate polycomb domains every cell cycle

**DOI:** 10.1101/2022.08.02.502476

**Authors:** Giovana M. B. Veronezi, Srinivas Ramachandran

## Abstract

Gene repression by the polycomb pathway is essential for proper development in metazoans. Polycomb domains, characterized by trimethylation of histone H3 (H3K27me3), carry the memory of repression through successive cell divisions, and hence, need to be maintained to counter dilution during replication of parental H3K27me3 with unmodified nascent H3. Yet, how locus-specific H3K27me3 is maintained through replication is unknown. To understand H3K27me3 recovery post-replication, we first defined nucleation sites within each polycomb domain in mouse embryonic stem cells. Then, to track H3K27me3 dynamics in unperturbed cells during replication, we developed CUT&Flow, which maps H3K27me3 domains as a function of the cell cycle stage. Using CUT&Flow, we show that post-replication rejuvenation of polycomb domains occurs by nucleation and spreading, using the same nucleation sites used during *de novo* domain formation. By using subunit-specific inhibitors of the PRC2 complex, we find that PRC2 need not rely on pre-existing H3K27me3 to target nucleation sites post-replication. The timing of nucleation relative to the cell cycle reflects the combination of replication timing and H3K27me3 deposition kinetics derived from *de novo* domain formation. Thus, competition between H3K27me3 deposition and nucleosome turnover drives both *de novo* domain formation and maintenance during every cell cycle.

## Introduction

Many chromatin states are cell-type specific and maintained through successive cell divisions. One such chromatin state is the trimethylation of histone H3 lysine 27 (H3K27me3), catalyzed by Polycomb Repressive Complex 2 (PRC2). The Polycomb and Trithorax groups (PcG and TrxG) of proteins are essential for proper cell fate specification during development^1^. Regions silenced by PcG proteins form broad domains of H3K27me3 on the genome^2,3^. These domains underlie the cell type-specific silencing by PcG, but the mechanisms of their maintenance despite dilution during replication remains unclear.

Several factors drive locus-specific deposition of H3K27me3. Core PRC2 consists of EZH2, SUZ12, RBAP46/48, and EED, with EZH2 being the catalytic subunit^4–6^. SUZ12 is the central scaffold and has been implicated in directing PRC2 activity genome-wide^7,8^. PRC2 spreading is thought to depend on its EED subunit. EED can specifically bind H3K27me3, which allosterically increases the catalytic activity of the EZH2 subunit^9^. Thus, EED can bind pre-existing H3K27me3 and stimulate EZH2 to modify a spatially adjacent H3 tail. Polycomb Repressive Complex 1 (PRC1)^10^ is an E3 ubiquitin ligase that mono-ubiquitinates H2A K119^11^. H2A ubiquitination can drive PRC2 targeting^12,13^. Finally, H3.3 deposition is important for PRC2 activity^14^, possibly because PRC2 nucleation sites may feature nucleosome turnover and have DNA accessibility^15^.

Studies in *Drosophila*, murine, and human cells have shown that two mechanisms lead to H3K27me3 deposition. In the first, specific DNA sequences act as nucleation sites for PRC2^16–19^. Early work in *Drosophila* identified sequences in Hox clusters that could silence transgenes in a cell type specific manner and these sequences were denoted as Polycomb Response Elements (PREs)^20,21^. PREs are thought to be direct binding sites of PRC2. Second, via its EED subunit, PRC2 can then use the pre-existing H3K27me3 to modify nearby unmodified histones through a read-write mechanism similar to DNA methyltransferases^9,22^. These two mechanisms have been succinctly described as “nucleation” and “spreading”^23^. Nucleation and spreading has been observed in a system where H3K27me3 domains were ablated and then reconstructed *de novo* in an inducible manner^22^.

DNA replication dilutes global H3K27me3 levels two-fold. Hence, H3K27me3 restoration must occur every cell cycle after DNA is replicated. Tracking H3K27me3 levels through the cell cycle in HeLa cells has shown that levels of this modification are restored predominantly in the G1 phase following replication^24^. However, in mammalian cells, it is unknown if nucleation drives the restoration of H3K27me3 post-replication or if the parental H3K27me3 redeposited across the domain^25,26^ drive methylation of the nascent H3. Technical challenges in tracking H3K27me3 dynamics across the cell cycle have limited understanding the mechanism of replenishment post-replication. With *de novo* domain formation after PRC2 ablation, the H3K27me3 levels start with a clean slate and go from almost zero to steady-state levels. In contrast, the change in H3K27me3 during the cell cycle is within a much smaller range of 50-100%^27^. Importantly, previous studies tracking H3K27me3 temporally post-replication have not asked if the nucleation and spreading model of *de novo* domain formation applies to H3K27me3 maintenance post-replication. Moreover, even if nucleation operated during the cell cycle, it is not known if nucleation sites used during *de novo* domain formation would also be used after replication.

To ask if nucleation occurs in unperturbed cells during the cell cycle, we developed a framework to identify nucleation sites based just on the intradomain kinetics of H3K27me3. The nucleation and spreading model predicts that, within a polycomb domain, the nucleation sites should have the fastest kinetics of increase in H3K27me3 enrichment. We first partitioned chromosomes into H3K27me3 domains and then identified nucleation sites for each of those domains. We then asked if nucleation occurs in unperturbed cells. To do this, we developed a new method, CUT&Flow, by coupling Cleavage Under Target and Tagmentation (CUT&Tag) with flow cytometry, to define H3K27me3 domains as a function of the cell cycle stage. With CUT&Flow, we observe nucleation at specific cell cycle stages, at the same sites that nucleate during *de novo* domain formation. The cell cycle stage when a site nucleates is determined by a combination of its replication timing and the site’s inherent H3K27me3 deposition kinetics. Strikingly, the nucleation sites are more sensitive to inhibition of EZH2 compared to inhibition of EED (which enables PRC2 to recognize pre-existing H3K27me3). The higher sensitivity of PRC2 to catalytic inhibition compared to allosteric inhibition confirms that PRC2 nucleation occurs post-replication. Thus, the opposing forces of PRC2 nucleation and DNA replication can explain the steady-state landscape of Polycomb domains genome-wide. Our results support the idea that a unified nucleation mechanism governs both *de novo* domain formation and rejuvenation of H3K27me3 domains every cell cycle.

## Results

### Large Polycomb domains have diverse intra-domain H3K27me3 kinetics

Because Polycomb domains are expected to be on the scale of kilobases, we reasoned that traditional analysis of H3K27me3 peaks would not be appropriate to analyze H3K27me3 recovery through the cell cycle. Genome-wide peak calling alone would mask the identification of nucleation and spreading occurring within a polycomb domain, as the positional information of H3K27me3 dynamics within individual Polycomb domains is lost. To overcome this limitation, we first devised a simple domain calling algorithm and applied it to murine embryonic stem cells (mESCs) using published epigenomic data (see Methods). We identified domains at various scales from 1 kb to greater than 100 kb (**Figure S1**). As expected, our algorithm identified large domains at Hox clusters, which are known to have high enrichment of H3K27me3 in mESCs (**Figure S2**). In a published study^8^, mESC were treated with an Ezh2 inhibitor for 10 days, which resulted in a reduction of H3K27me1, H3K27me2, and H3K27me3 to levels undetectable by immunoblots. The inhibitor was then washed out and recovery of H3K27 methylation states was tracked by immunoblot and spike-in normalized ChIP-seq 4-, 8-, 16-, 24-, 48-, and 96-hours post-washout. We used this timed ChIP-seq dataset and our identified domains in mESCs to develop a framework for tracking intradomain temporal dynamics of H3K27me3.

We reasoned that the absolute enrichment of H3K27me3 at steady state could obscure differences in kinetics between nucleation sites and the rest of the domain for several reasons. First, earlier studies have shown that nucleation sites feature high nucleosome turnover^15^. Second, H3.3 incorporation (another measure of turnover) is essential for the maintenance of Polycomb domains^14^, and *in vitro*, PRC2 binding itself might be stimulated by the presence of a nucleosome-free region^28^. Additional factors such as nucleosome density^29^ and H3.3-mediated deposition of the histone variant H2A.Z^30^ also positively regulate H3K27me3 deposition. Thus, to remove the effect of absolute H3K27me3 levels on observed kinetics, we plotted all time-points post-drug washout normalized to the final time-point (96-hour) ChIP signal. We observed a range of H3K27me3 recovery kinetics once we normalized 25 bp non-overlapping segments within a domain to their levels at 96 hours post-washout, exemplified by the Polycomb domain covering the *Hoxa* locus (**Figure 1A**). We next performed k-means clustering (k=7) of H3K27me3 recovery patterns at the normalized segments within each domain (**Figure 1B**). To order the clusters based on the speed of their recovery, we defined T_40_ as the time taken to reach 40% of steady-state H3K27me3 levels (**Figure 1C**). This was chosen because 40% represented a level at which there was a steady increase in H3K27me3 levels across the domain. T_40_ enabled automated ordering of the clusters from fastest to slowest recovery (**Figure 1B, C**). Importantly, the normalization did not affect the order of the clusters but enabled automated ordering of clusters (**Figure S3A-D**).

**Figure 1.**
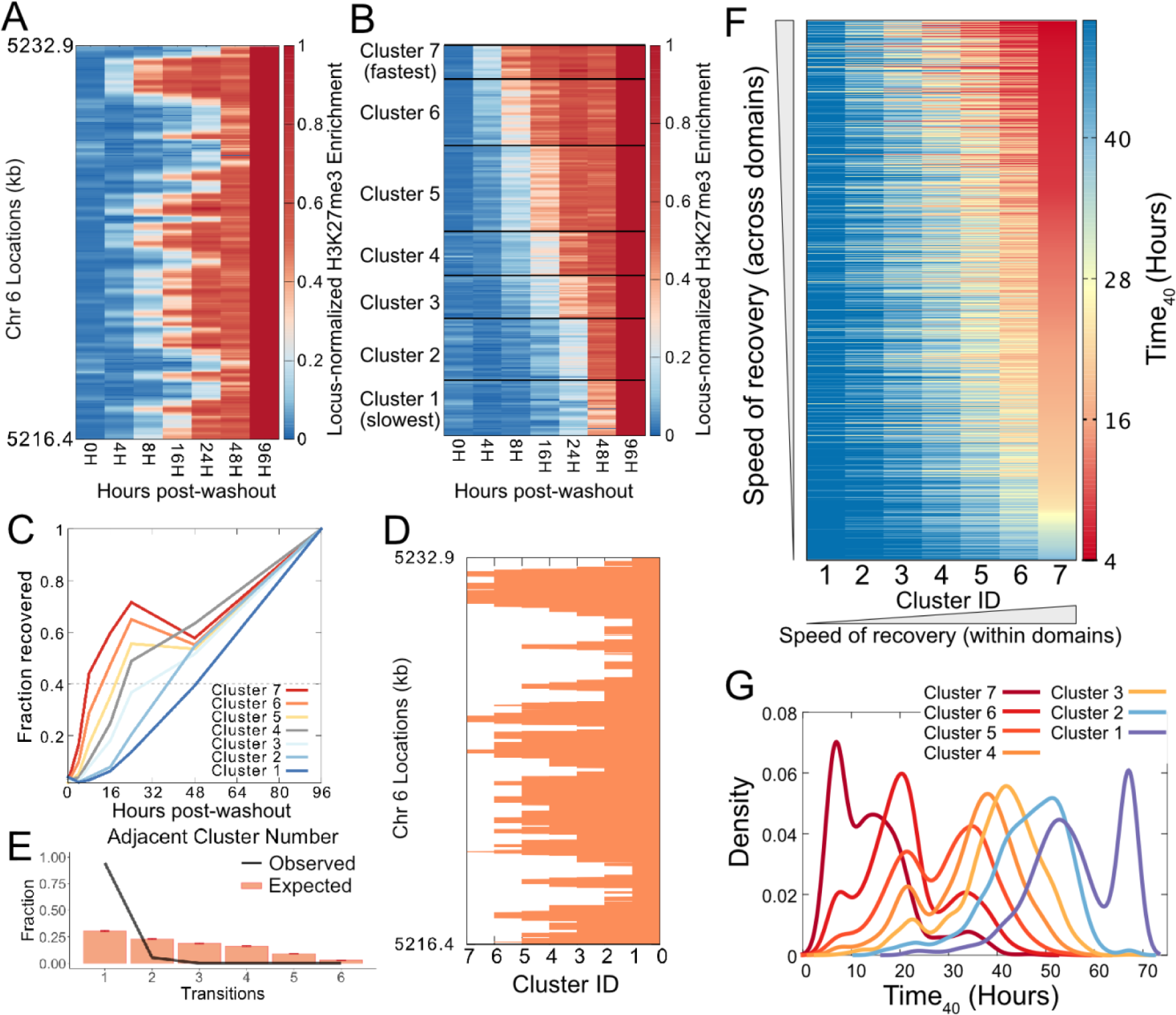
Intradomain kinetics of H3K27me3 recovery. **A**) Heatmap of H3K27me3 ChIP enrichment at time points post-washout of Ezh2 inhibitor (x axis) normalized to the ChIP enrichment at 96 hours post-washout. Each horizontal line of the heatmap represents 25 bp step of the genomic location on the Hoxa domain. **B**) Heatmap of H3K27me3 ChIP enrichment at time points post-washout of Ezh2 inhibitor (x axis) after performing k-means clustering with 7 clusters. Cluster 7 shows the fastest recovery of H3K27me3 post-washout, and cluster 1 shows the slowest recovery. **C**) Average H3K27me3 ChIP enrichment of each of the 7 clusters obtained by k-means clustering as a function of time post-washout. **D**) The k-means cluster IDs (7 is the fastest recovering cluster post-washout, 1 is the slowest) are plotted as a function of genomic location for the Hoxa domain. **E**) The observed transitions in cluster IDs that occur in adjacent genomic locations (in 25 bp steps) in the Hoxa domain is plotted as a black line. The expected transitions due to random recovery (instead of nucleation and spreading) is modeled by shuffling the cluster ID and the distribution is plotted as orange bars. **F)** Heatmap of T_40_ of each cluster in 1362 domains genome-wide. The clusters are obtained by k-means clustering performed on H3K27me3 ChIP values over time-points of recovery as shown in (**B**). For each domain, the clusters are ordered from slowest to fastest. **G**) Histogram of T_40_ for each cluster across domains shown as a heatmap.

The nucleation and spreading model posits the presence of nucleation sites which regain H3K27me3 first due to direct DNA binding of PRC2, and subsequently the other sites in the domain regain H3K27me3 due to spreading^23^. A consequence of this model is that adjacent sites will have similar kinetics and that H3K27me3 restoration kinetics will gradually change along the domain: in other words, a slow site will not be adjacent to a fast site. To ask if this was the case, we mapped the kinetic clusters back onto the domain and observed a “peak and valleys” distribution of cluster id that would be expected for nucleation and spreading (**Figure 1D, Figure S3E**). Even though the k-means clustering did not use positional information, boundaries between kinetic clusters were strongly enriched for transitions to successive clusters, indicating that there was a stepwise change in the rate of H3K27me3 recovery (**Figure 1D, E**). The stepwise change could not be reproduced when cluster numbers within the domain were shuffled, indicating that this was due to the non-random distribution of kinetic clusters (**Figure 1E**). This strongly suggests that the spreading of PRC2 from nucleation sites could explain the observed kinetics of H3K27me3 post drug washout. Even though H3K27me3 patterns return to pre-treatment, steady-state distribution ∼96 hours after drug washout, our analysis shows that the kinetics of recovery is not uniform.

### Polycomb domains are independent units for PRC2 kinetics

We hypothesized that the kinetic cluster within each domain with the fastest H3K27me3 recovery represents nucleation sites for PRC2. Interestingly, when we compared the H3K27me3 recovery at large domains defined at Hoxa and Hoxd gene clusters, we observed a significant difference in their kinetics: the average T_40_ of the fastest kinetic group of the Hoxa domain was ∼7 h 23 m, whereas that of the fastest kinetic group in the Hoxd domain was only 5 h. In other words, even though both the domains feature characteristics of nucleation and spreading, Hoxa nucleation is 32% slower than Hoxd. This result led us to ask how the T_40_ of nucleation sites varies across domains genome-wide. We chose 1362 domains that were common across different datasets and at least 10 kb in length. Within each domain, we performed k-means clustering of H3K27me3 spike-in ChIP across the time points, normalized to the 96-hour time point, then sorted the kinetic clusters from slowest to fastest based on each cluster’s average T_40_. We observed a broad diversity in H3K27me3 recovery genome-wide, with nucleation sites having T_40_ values ranging from 3h 19m to nearly 48 hours (**Figure 1F**). This wide range in recovery times across domains is reflected in the highly overlapping distribution of T_40_ values for each cluster (**Figure 1G**). Thus, each domain has its characteristic nucleation rate, presumably due to differences in PRC2 recruitment strength at their nucleation sites. This analysis also demonstrates an important point that nucleation and spreading within each domain might not be apparent if recovery kinetics at H3K27me3 peaks were compared across whole chromosomes instead of analyzing intradomain kinetics for individual domains. When compared across chromosomes, we would only identify the fastest recovering domains rather than how each domain itself recovers. For nucleation sites, shorter T_40_ values would denote stronger PRC2 nucleation. At the Hoxd locus, two of our predicted nucleation sites overlapped with previously characterized nucleation sites: Hoxd11-12^18^, and Evx^22^, consistent with intradomain kinetics being able to identify nucleation sites. We also validated the intradomain nucleation sites we identified using published data from EED cage mutants. EED binds H3K27me3 through an aromatic cage^9^. Mutations of residues in the aromatic cage result in loss of EED binding to H3K7me3. EED’s loss of binding to H3K27me3 results in PRC2 unable to spread^22^. Oksuz *et al.*^22^ introduced these mutations in mESCs and then performed H3K27me3 ChIP. In the cage mutants, the enrichment of H3K27me3 at the nucleation sites we identified based on recovery kinetics was even more exaggerated compared to WT, leading to an apparent increase in nucleation strength with these mutants (**Figure S4A**). In summary, by considering each domain individually, we not only found that each domain has its unique nucleation rate, but also identified putative nucleation sites for each of those H3K27me3 domains.

### CUT&Flow profiles Polycomb domains across the cell cycle

Nucleation is the obvious means to establish a domain *de novo*. However, post-replication, domains still have 50% of parental H3K27me3 that can attract PRC2 via its EED subunit. Thus, given the abundance of pre-existing H3K27me3, it is not clear if PRC2 nucleation would still significantly occur post-replication as polycomb domains are replenished. We reasoned that, if nucleation occurred post-replication, the enrichment of H3K27me3 at nucleation sites relative to spreading sites would be highest at the early stages of rejuvenation of a domain. To track polycomb domain dynamics across the cell cycle in unperturbed cells, we hypothesized that cells in different stages of the cell cycle would feature H3K27me3 domains with different extents of nucleation and spreading. To capture H3K27me3 domains at various stages of the cell cycle, we combined H3K27me3 profiling by Cleavage Under Targets and Tagmentation (CUT&Tag)^31^ with flow-sorting to fractionate cells at different stages of the cell cycle. The technique, which we term CUT&Flow, involves similar steps to CUT&Tag until tagmentation: isolation of nuclei, binding of primary and secondary antibodies, binding of pA-Tn5, and tagmentation. The nuclei are then labeled with propidium iodide and sorted into 5 gates based on DNA content: G1, early S, mid S, late S, and G2/M (**Figure 2A, B**). Finally, the sorted nuclei are lysed, and PCR is performed to obtain sequencing libraries, each one representing H3K27me3 profiles of cells at a specific stage of the cell cycle.

**Figure 2.**
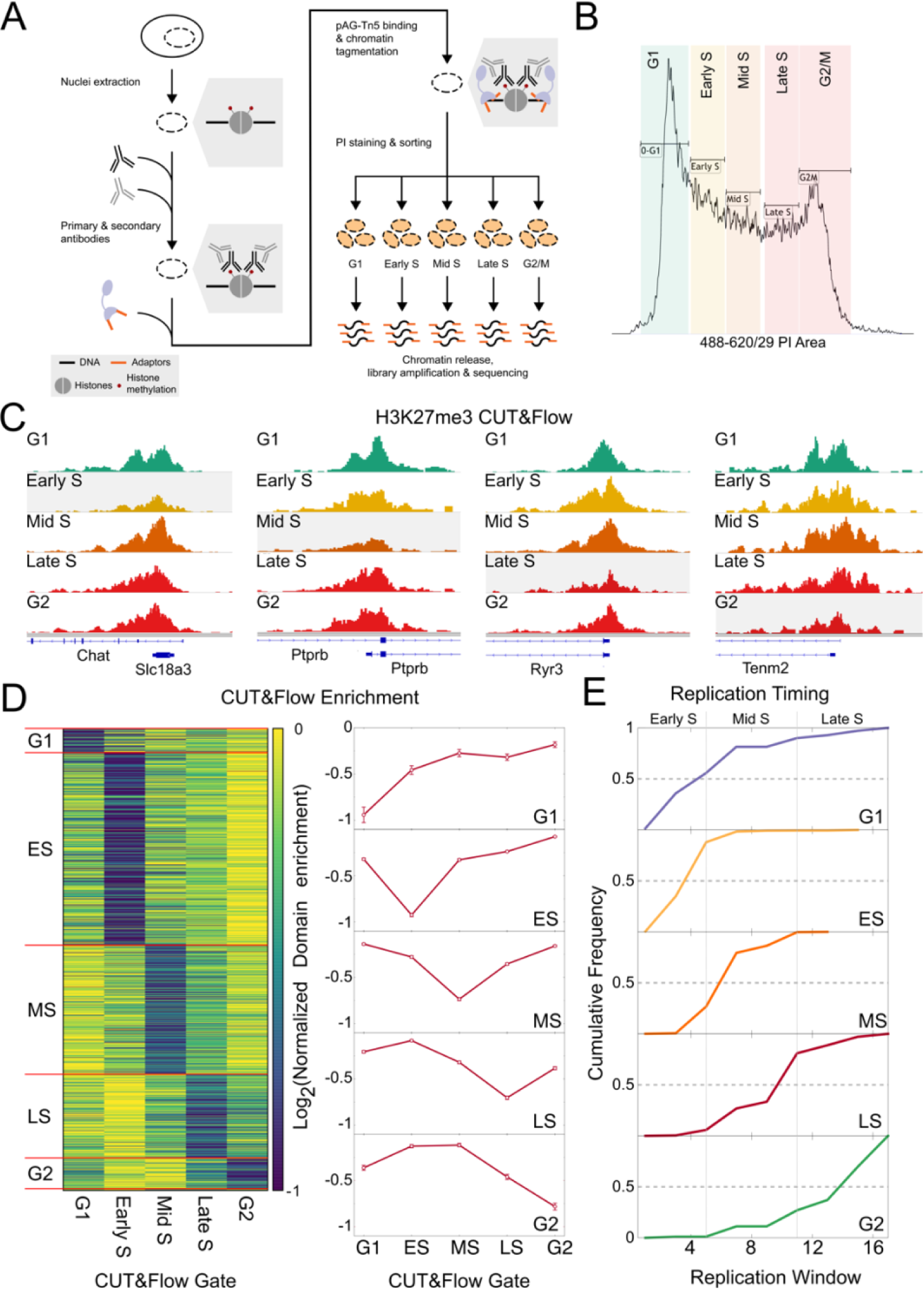
CUT&Flow captures cell cycle stage-specific chromatin profiles. **A**) Schematic of the CUT&Flow protocol. **B**) Distribution of Propidium Iodide fluorescence of nuclei. The 5 gates to segregate nuclei based on stage of cell cycle are shown. **C**) Examples of domains where the normalized enrichment of H3K27me3 drops significantly in only one of the CUT&Flow gates. The gate with significant drop is shaded gray. The genomic regions displayed are chr14:32,447,403-32,471,747 (first from left), chr10:116,295,946-116,306,843 (second from left), chr2:113,204,914-113,225,200 (second from right), and chr11:37,224,873-37,244,919 (first from right). **D**) Heatmap (left) showing classification of domains based on the CUT&Flow gate which shows the lowest H3K27me3 enrichment. The enrichments for each domain are normalized to the gate with highest H3K27me3. Mean and S.E.M. for each classification from the heatmap are shown as a line plot (right). **E**) Cumulative histogram of the S-phase window (1-16) in which each group of domains sites (defined in (**D**)) replicate most often. Domains that have reduced enrichment in G1 and Early S have earliest replication timing, followed by Mid S, Late S, and then G2.

To ask if CUT&Flow profiles H3K27me3 dynamics across the cell cycle, we first took advantage of the fact that DNA replication dilutes H3K27me3 levels. We asked if H3K27me3 enrichment was diluted at specific polycomb domains in the different cell cycle gates of CUT&Flow. Notably, just by visual inspection, individual domains showed a significant drop of H3K27me3 enrichment at just one cell cycle gate followed by recovery in the next gates (**Figure 2C**). Extending this analysis genome-wide, we assigned a CUT&Flow cell cycle gate to each domain based on when that domain featured a sudden drop in H3K27me3. We were able to assign a cell cycle gate to most of the domains (**Figure 2D**). The highest level of H3K27me3 for a domain occurred at the third cell cycle gate downstream of the gate at which dilution occurred, suggesting dilution followed by recovery of H3K27me3 due to replication.

We observed the dilution of H3K27me3 to be around one-half fold (**Figure 2D, right**), consistent with the expected level of dilution caused by replication. If the dilution we observe in CUT&Flow gates occurred due to replication, then the replication timing of the domains should correlate with the timing of the dilution. To determine the replication timing of the Polycomb domains, we analyzed published Repli-Seq data for mESC^32^, where cells were sorted into 16 S-phase fractions after BrdU labeling. Then, BrdU IP was performed followed by sequencing. The S-phase fraction with the highest Repli-Seq signal for a genomic region represents the predominant replication timing of that region. The domains that underwent dilution in G1 and early S replicated earliest, followed by mid S, late S, and finally G2 (**Figure 2E**). Thus, replication timing strongly correlated with the timing of H3K27me3 dilution mapped by CUT&Flow and leads us to conclude that CUT&Flow indeed captures H3K27me3 dynamics across the cell cycle in unperturbed cells.

### H3K27me3 domains feature nucleation post-replication

To observe dynamics of H3K27me3 at nucleation sites across the cell cycle, we calculated the ratio of H3K27me3 enrichment at nucleation sites to the enrichment at spreading sites for the different CUT&Flow gates. We first plotted the spreading sites-normalized H3K27me3 profiles from G1, Early S, Mid S, Late S, and G2 at the Hoxd cluster and observed robust H3K27me3 domains with similar boundaries to steady-state profiles (**Figure 3A**). Interestingly, we observed an increase in H3K27me3 enrichment at sites corresponding to nucleation sites and sites adjacent to nucleation sites (kinetic clusters 6 and 7; kinetic clusters defined in **Figure 1**) in the Mid S dataset compared to the other datasets. We calculated the change in enrichment over the cell cycle for sites in each kinetic cluster normalized by enrichment in spreading sites (kinetic clusters 1 and 2). Clusters 3 and 4 did not fluctuate much over the cell cycle, but clusters 5, 6, and 7 showed a peak in Mid S, with cluster 7 having the strongest increase in Mid S (**Figure 3B**). From published Repli-Seq data for mESC^32^, we observed the Hoxd locus to predominantly replicate in Mid S (**Figure 3C**). Thus, the spike in enrichment in nucleation sites occurred close to when the domain was replicated, pointing to rapid nucleation following replication. We next looked at the nucleation sites in Hoxa and Hoxc clusters and observed a spike in enrichment in Early S and Mid S respectively (**Figure 3D**). When we plotted the Repli-Seq data for these two clusters, we observed both clusters to replicate in Early S (**Figure 3E**). Thus, Hoxa and Hoxd clusters feature a spike in nucleation close to when their respective polycomb domains replicate, but there is a delay in the spike for the Hoxc cluster compared to its replication timing. The delay in Hoxc could be due to slower nucleation kinetics in Hoxc compared to Hoxa. We asked if the *de novo* nucleation data also showed Hoxc to have slower nucleation than Hoxa. We have a measure of *de novo* PRC2 nucleation speed as T_40_. Indeed, the T_40_ of Hoxc is significantly longer than Hoxa, demonstrating a correlation between *de novo* nucleation kinetics and post-replication spike in H3K27me3 at nucleation sites (**Figure 3F**). Our results suggest that there is cell cycle-dependent nucleation in the Hoxa, Hoxc, and Hoxd polycomb domains and this nucleation appears to be coupled to the replication of the respective domains.

**Figure 3.**
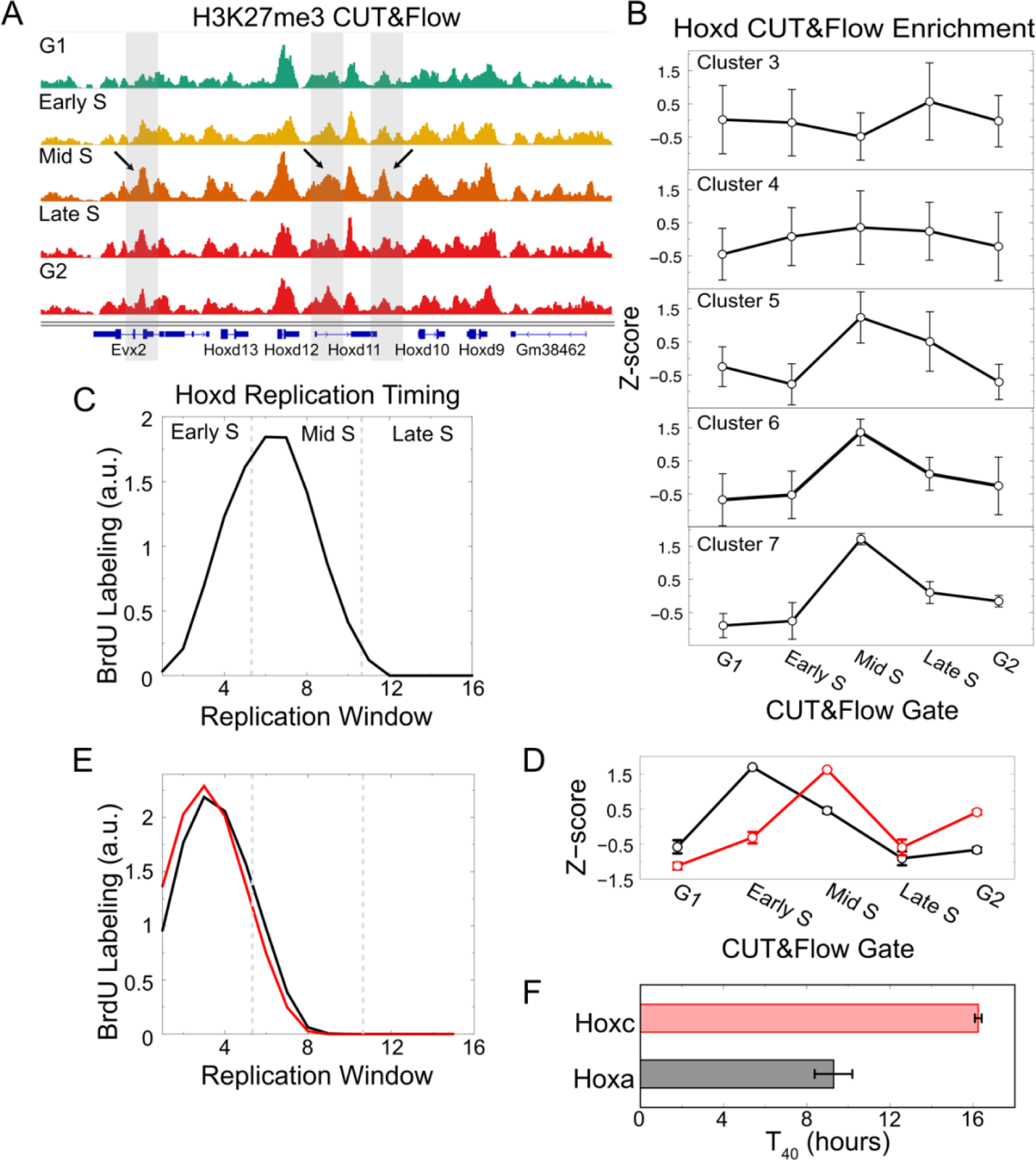
CUT&Flow captures cell cycle stage-specific nucleation site dynamics at Hox loci. **A**) H3K27me3 profiles at the Hoxd locus from each CUT&Flow gate. The regions shaded in gray are nucleation sites. The arrows point to peaks with increased enrichment in the mid S. **B**) Quantification of enrichment of regions in each kinetic cluster (defined in Figures 2 and 3) normalized to enrichment of Clusters 1 and 2 shows a significant increase for Clusters 5, 6, and 7 at Mid S. **C**) Repli-seq enrichment of the Hoxd locus from published data indicates that the locus replicates in Mid S. **D**) Similar to (**B**) for nucleation sites in Hoxc and Hoxa clusters. **E**) Similar to (**C**) for Hoxc and Hoxa clusters. **F**) Bar plot showing mean ± S.E.M. of the T_40_ for nucleation sites of Hoxc and Hoxa clusters.

### PRC2 kinetics predict cell cycle-dependent nucleation

To examine cell cycle dynamics of nucleation sites genome-wide, we calculated the ratio of H3K27me3 enrichment at each nucleation site to the enrichment at their corresponding spreading sites for the different CUT&Flow gates. Based on the nucleation behavior of the Hoxd locus, this ratio is predicted to spike at a specific cell cycle stage when the domain nucleates. We analyzed cell cycle stage-specific enrichment of H3K27me3 at nucleation and spreading sites that were defined based on *de novo* domain formation (in **Figure 1**). For most of the domains, we found the nucleation site enrichment increased dramatically in one phase of the cell cycle and then declined (**Figure 4A**). These strong cell cycle stage-specific events were marked by H3K27me3 levels at least 1 Z-score higher than other cell cycle stages, followed by a decrease presumably as H3K27me3 spread through the domain (**Figure 4A**). The spreading sites showed a decrease in Z-scores compared to the nucleation sites, and this decrease correlated with their distance from the nucleation site. Thus, the strongest gain in H3K27me3 was at the nucleation sites at specific cell cycle stages, followed by sites adjacent to the nucleation sites, and so on (**Figure 4B**). To confirm that the spike occurred on newly replicated chromatin, we analyzed an independent, published dataset generated using ChOR-seq^33^. For H3K27me3 ChOR-seq, cells are labeled with EdU followed by immunoprecipitation of H3K27me3-containing nucleosomes, isolation of EdU-labeled DNA and sequencing. Strikingly, we observe a transient increase in H3K27me3 enrichment at the same nucleation sites in ChOR-seq data 3 hours post-replication (**Figure S4B, C**), independently validating the nucleation post-replication observed with CUT&Flow. Our results suggest that the post-replication rejuvenation of polycomb domains occurs by nucleation and spreading, via the same nucleation sites used during *de novo* domain formation.

**Figure 4.**
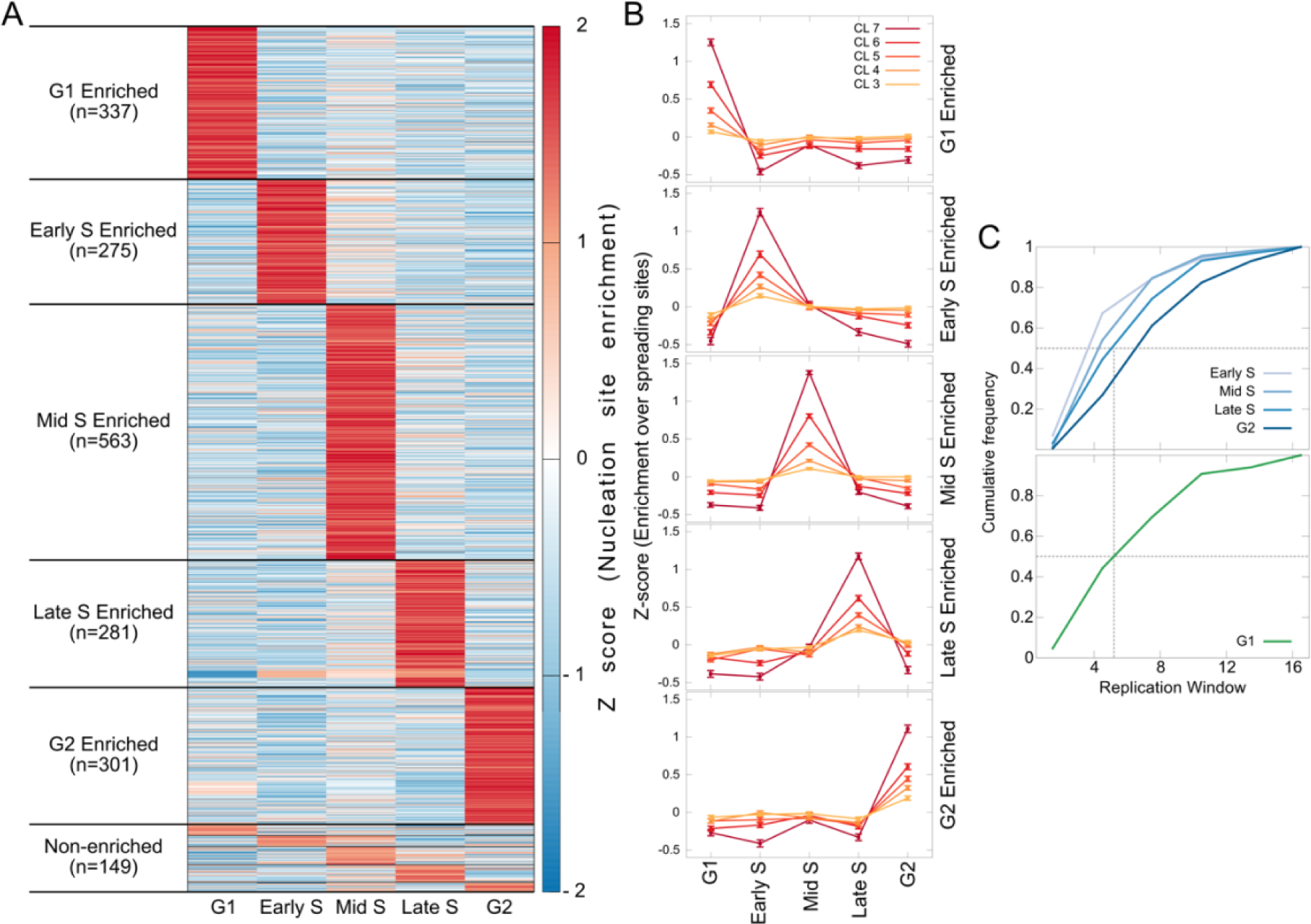
CUT&Flow shows H3K27me3 nucleation post-replication. **A**) Z-score of nucleation site enrichment over spreading sites across the cell cycle plotted as a heatmap. Nucleation sites are arranged based on the stage of the cell cycle that they spike in. **B**) For domains that feature spike in nucleation at each stage of cell cycle, the enrichment of H3K27me3 at clusters 3-7 over the enrichment at spreading sites (clusters 1,2) plotted as Z-scores. **C**) Cumulative distribution of the S-phase window (1-16) in which each group of nucleation sites (defined in (**A**)) replicate most often. Nucleation sites that spike in Early S have earliest replication timing, followed by Mid S, Late S, and G2 (top). G1 distribution is at the middle of the spread (bottom).

If these nucleation events occur post-replication, the replication timing of the domains that spike in early S is expected to be earlier than those of mid S, and so on, with the domains that spike in late S and G2 having the latest replication times. For each nucleation site, we determined the peak of the Repli-Seq signal. We observed the cumulative distribution of replication peak to be leftmost for those nucleation sites whose enrichment spikes in early S, followed by mid S, late S, and G2 (**Figure 4C, top**). Nucleation sites that spiked in G1 had replication timing similar to those domains that spiked in late S (**Figure 4C, bottom**). The correlation between the timing of H3K27me3 nucleation events determined by CUT&Flow and replication timing suggests that there is strong coordination between these processes. The fact that nucleation occurs immediately following replication points to rapid H3K27me3 deposition coupled with replication at nucleation sites in contrast to the slower global restoration of total H3K27me3^24^. Ezh2 and Jarid2 are enriched on chromatin in mESCs in S and G2 phases^34^, which agrees well with our observation of rapid nucleation coupled to replication. In summary, we observe a spike in H3K27me3 at nucleation sites of each domain, the timing of which correlates with the timing of the replication of the domain. Thus, PRC2 acts first at the same nucleation sites within domains for both *de novo* domain formation and restoration post-replication, implying that an intrinsic feature of these sites might drive PRC2 nucleation.

### Nucleation site chromatin properties identify distinct classes of Polycomb domains

To understand what mediated nucleation, we considered different molecular components that are important for PRC2 activity. CpG islands and bivalent promoters have been linked with PRC2 nucleation^8,22,35,36^. Jarid2 and PCL2 are DNA binding proteins that are auxiliary subunits of PRC2. Based on the well-characterized roles of Jarid2, PCL2, Ezh2, Suz12, H2AK119 ubiquitination (uH2A), chromatin accessibility, and H3.3 incorporation, we asked if these features are enriched at nucleation sites relative to sites with slow kinetics of H3K27me3 recovery. We observed significant enrichment of all these features at putative nucleation sites compared to the slowest sites in each domain, with the PRC2 components showing strongest enrichment (**Figure 5A**).

**Figure 5.**
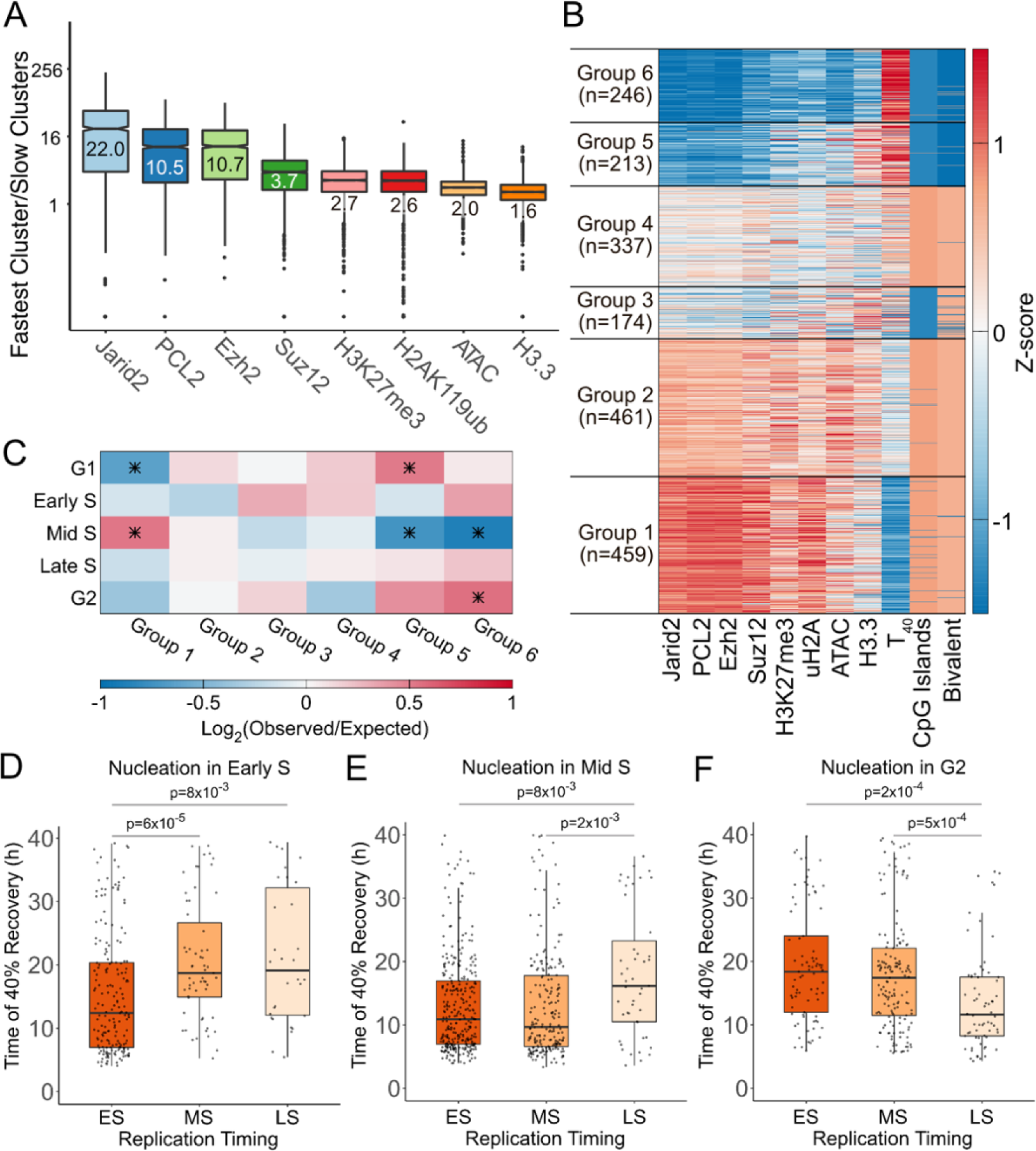
Chromatin properties at nucleation sites. **A**) Ratio of enrichment of different components/chromatin features at nucleation sites to the enrichment at spreading sites for all domains shown as boxplots. **B**) Groups obtained by k-means clustering of domains based on the z-score of features shown as a heatmap. **C**) Overlap of domains based on cell cycle stage-specific nucleation and groups based on chromatin features from (**B**). Overlaps that are significant by hypergeometric test (p<0.05, fdr<0.1) are denoted by asterisk. **D**) Distribution of the recovery kinetics of nucleation sites (measured by T_40_) plotted for domains separated by the window of replication, for domains that feature H3K27me3 nucleation spike in Early S. **E**) Same as (**D**) for domains that feature H3K27me3 nucleation spike in Mid S. **F**) Same as (**D**) for domains that feature H3K27me3 nucleation spike in G2.

We then asked if all these components are enriched to the same extent at nucleation sites across domains. We performed k-means clustering of the Z-scores of enrichments of these features at nucleation sites across domains to obtain six groups of domains. The presence/absence of bivalent promoters and CpG islands, and T_40_ at nucleation sites were then added in the same order as the identified groups of domains. Finally, we ordered the groups by T_40_ (**Figure 5B**). We observed that the PRC2 components, H3K27me3, uH2A, and ATAC clustered together and had the highest enrichment at domains with the fastest nucleation. H3.3 was enriched in groups 2, 3, and 5, with a pattern that was distinct from Polycomb components. Group 5, with the highest levels of H3.3, also featured slower nucleation. This analysis points to three general classes of nucleation sites: fast nucleation sites that feature strong binding of PRC1 and PRC2 at steady state; slower nucleation sites that feature H3.3; and slowest nucleation sites that had low levels of all components surveyed. Interestingly, CpG islands and bivalent promoters overlap with groups 1, 2, and 4, underlining the fact that sequence only partially explains PRC2 kinetics. Transcription start sites overlapping with the domains in group 1 were enriched for neuronal development/developmental genes and transcription factors, whereas group 2 was enriched in genes encoding membrane proteins (**Table S3**). In summary, our analysis suggests that the components that drive PRC2 activity are enriched at nucleation sites relative to spreading sites at steady state and the extent of enrichment at steady state correlates with the speed of H3K27me3 deposition post drug washout.

We next asked if the cell cycle stage-specific nucleation of H3K27me3 correlated to the chromatin features of the domain. To do this, we intersected the cell cycle stage of the nucleation of a domain with the groups defining chromatin/sequence states of the domain. We observed a high overlap that was statistically significant between Group 1 and the nucleation spike in mid S (**Figure 5C**). We observed depletion of Group 1 and enrichment of Group 5 domains with nucleation spike in G1. Finally, we observed an enrichment of Group 6 domains with a nucleation spike in G2. Group 1 has the fastest kinetics in *de novo* domain formation and Group 6 has the slowest kinetics (**Figure 5B**). In other words, fast nucleation sites nucleate in Mid S, and slow nucleation sites nucleate in G1 and G2, suggesting a correlation between nucleation kinetics in *de novo* domain formation and nucleation during the cell cycle (**Figure 5C**). This led us to ask if PRC2 nucleation kinetics was coupled to replication timing. We have a measure of *de novo* PRC2 kinetics as the time to 40% recovery after drug washout (T_40_). If a nucleation site displays a large gap between replication and nucleation during the cell cycle, for example, if a site replicates in early S but nucleates only in G1, that could be due to slow PRC2 nucleation kinetics at that site. To ask if this is the case, we first divided nucleation sites that spike in each phase of the cell cycle into their replication timing bin based on Repli-Seq: early S, mid S, and late S. We then plotted the T_40_ for each replication timing bin. For domains nucleating in early S, we observe a majority of them replicated in early S as expected (**Figure 5D**). The domains that replicated in Early S displayed significantly faster kinetics than those that replicated in mid and late S. The mid and late S replicating domains that nucleate in early S surprisingly have replication timing that would be after nucleation in early S. Thus, these sites seem to nucleate pre-replication, presumably to account for slower nucleation kinetics. We observe a similar phenomenon for domains that nucleate in mid S – the domains that replicate in early and mid S have significantly faster kinetics than those that replicate in late S (**Figure 5E**). Finally, for domains that nucleate in G2, those that replicate in late S have significantly faster kinetics than those that replicate in early and mid S (**Figure 5F**). Thus, H3K27me3 kinetics during *de novo* domain formation correlates with the time between replication and nucleation during the cell cycle. Our results suggest that a balance of locus-specific nucleation strengths and replication timing result in the genome-wide H3K27me3 landscape observed at steady state.

### CUT&Flow captures changes in PRC2 kinetics at short timescales

We next asked if the nucleation of H3K27me3 coupled with replication is due to PRC2 activity by performing CUT&Flow in the presence of well-characterized PRC2 inhibitors. Tazemetostat (EPZ-6438) is a potent, selective Ezh2 inhibitor^37^. EED226 is an allosteric inhibitor of PRC2, competing with H3K27me3 to bind EED^38^. Over 24 hours, we observed a similar decrease in H3K27me3 in mESC in the presence of EPZ-6438 or EED226 (**Figure S5**). Most previous studies treat cells with these inhibitors for multiple cell cycles to detect changes in H3K27me3^8,37^. Since CUT&Flow can capture transient nucleation events coupled to replication, we hypothesized that we could observe the effects of PRC2 inhibition in the timescale of hours before drastic changes in global H3K27me3 due to replicative dilution. With CUT&Flow, we could ask two questions with these inhibitors that were not possible before: i) what is the immediate effect of inhibiting PRC2 catalytic activity on H3K27me3 domains post-replication? ii) what is the effect of PRC2 inhibition on nucleation post-replication?

Treatment of mESC with EPZ-6438 or EED226 for four hours reduced H3K27me3 levels by 25-35%, while eight-hour treatments reduced H3K27me3 levels by 50% by flow cytometry. Thus, both inhibitors had similar effects on global levels of H3K27me3 in mESCs, even though they act via different mechanisms (**Figure 6A**, left panel). We next asked if treatment with the inhibitors affected the dilution and recovery dynamics of total H3K27me3 through the cell cycle using flow cytometry. In untreated cells, we observed a decrease in H3K27me3 global levels relative to G1 as cells enter early S, reaching the lowest levels in late S and recovering in G2 (**Figure 6A**; middle and right panel). Interestingly, the H3K27me3 dilution and recovery dynamics in mESCs treated with inhibitors recapitulated the pattern observed for untreated cells for both EPZ-6438 and EED226 treatments when normalized to G1 levels (**Figure 6A**; middle and right panel). Thus, although inhibitor treatments reduce levels of H3K27me3, the dynamics of global H3K27me3 dilution and recovery throughout the cell cycle are similar. We next asked how EPZ-6438 and EED226 treatments affected H3K27me3 recovery post-replication in a locus-specific manner.

**Figure 6.**
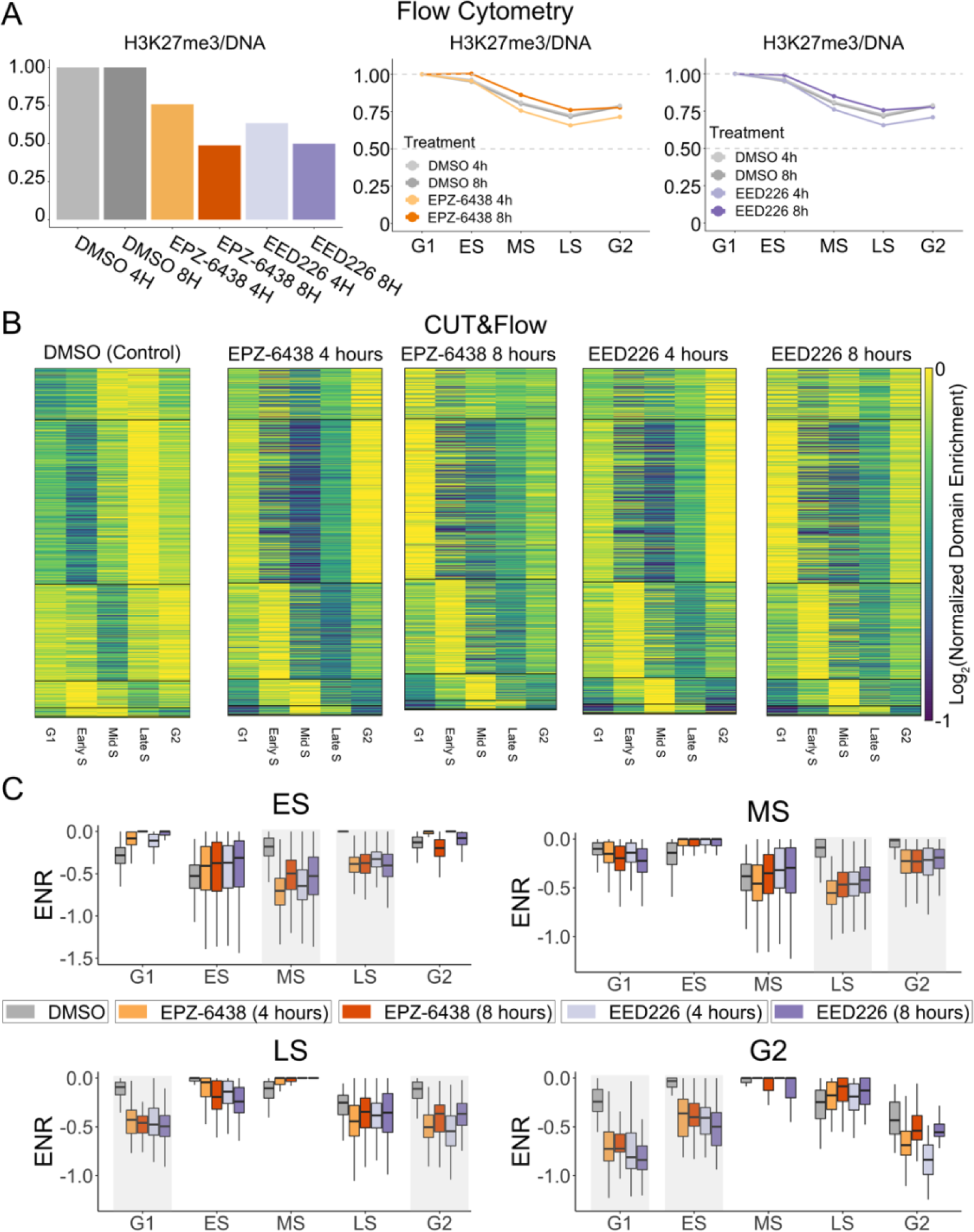
CUT&Flow shows delayed recovery of H3K27me3 post-replication upon PRC2 inhibition. **A**) Median H3K27me3 levels normalized to control (DMSO) samples plotted for 4h and 8h treatments with DMSO, EPZ-6438 and EED226 (left). Total H3K27me3 and DNA levels were determined by flow cytometry. H3K27me3 levels across the cell cycle were determined by gating for DNA content (middle and right). Median H3K27me3 levels were normalized so that G1 levels were 1 across treatments (middle and right). **B**) Heatmap (left, DMSO control) showing classification of domains based on the CUT&Flow gate which shows the lowest H3K27me3 enrichment. The enrichments for each domain are normalized to the gate with highest H3K27me3. The same classification was assigned to all treatments with PRC2 inhibitors and plotted similarly to control. **C**) Quantification of the heatmaps in (**B**) shown as boxplots for each assigned replication gate. The gray shaded boxes are the two gates that follow the assigned replication gate. With both inhibitors, it is seen that the H3K27me3 levels are lower compared to control in the shaded gates.

We performed H3K27me3 CUT&Flow after 4 and 8 hours of treatment of mESC with either EPZ-6438 or EED226. We first assigned a CUT&Flow gate to every domain based on when the domain has a decrease in H3K27me3 levels in our control dataset (similar to the analysis presented in **Figure 2**). We then plotted the domain enrichments of H3K27me3 across CUT&Flow gates for all the treatments in the same order as that of control (**Figure 6B**). We observed similar patterns for EPZ-6438 and EED226 at four and eight hours of treatment. Though we observed the dilution in the samples treated with the inhibitors at a similar gate as DMSO, for all inhibitor treatments, the lower levels of H3K27me3 persisted for two more gates, showing impaired recovery of H3K27me3 after replication (**Figure 6B, C**). Thus, even with treatment as short as 4 hours and a reduction in global H3K27me3 levels by just 25%, CUT&Flow captures a drastic decrease in PRC2 activity post-replication. These experiments thus validate our earlier conclusions that CUT&Flow captures H3K27me3 dynamics across replication dilution and recovery.

### EED226 has a weaker effect at nucleation sites compared to EPZ-6438

We next determined the effects of both inhibitors at nucleation sites. PRC2 nucleation depends on EZH2 activity much more than EED activity. Therefore, EPZ-6438 (which blocks EZH2 catalytic activity directly) should have a much bigger impact on nucleation compared to EED226 (which blocks EED subunit from binding methylated lysines). Thus, these two inhibitors’ distinct mechanisms of action allow us to uncouple PRC2 recognition of pre-existing H3K27me3 from direct recruitment of PRC2 to nucleation sites. We first calculated the enrichment of H3K27me3 at each nucleation site relative to its spreading sites (similar to the analysis presented in **Figures 3 and 4**). We then asked how this enrichment changed in the presence of the two inhibitors by calculating the log2 ratio of the maximal enrichment at nucleation sites for inhibitor treatment compared to control. At 4-, and 8-hour treatments, we observed a lower log2 ratio across nucleation sites for EPZ-6438 than EED226 (**Figure 7A, B**; most nucleation sites are above the diagonal). Globally, EPZ-6438 had a decrease in enrichment at nucleation sites compared to the control, and this decrease was significantly higher compared to EED226 (**Figure 7C**). Thus, catalytic inhibition of PRC2, which will affect nucleation and spreading, significantly blunts the post-replication spike in H3K27me3 at nucleation sites. In contrast, allosteric inhibition of PRC2, which will affect spreading much more than nucleation, has a minor effect on the post-replication spike in H3K27me3 at nucleation sites. The differential effects of the catalytic and allosteric inhibitors on H3K27me3 at nucleation sites post-replication highlight PRC2 nucleation that occurs post-replication at specific sites inside polycomb domains. The fact that the effects are similar at both four and eight hours of treatment with EPZ-6438 suggests that the spike at nucleation sites is an early event post-replication, confirming the correlation between replication timing and nucleation observed in unperturbed cells and the spike seen three hours post-replication in ChOR-seq data (**Figure S4**). The similar effects at four and eight hours of treatment, where global levels of H3K27me3 are lower by 25% and 50%, respectively, also shows that the enrichment at nucleation sites post-replication does not depend on pre-existing H3K27me3 levels. In summary, by separating allosteric recognition and catalysis, we establish that PRC2 predominantly acts at nucleation sites post-replication and spreads from there to restore H3K27me3 levels.

**Figure 7.**
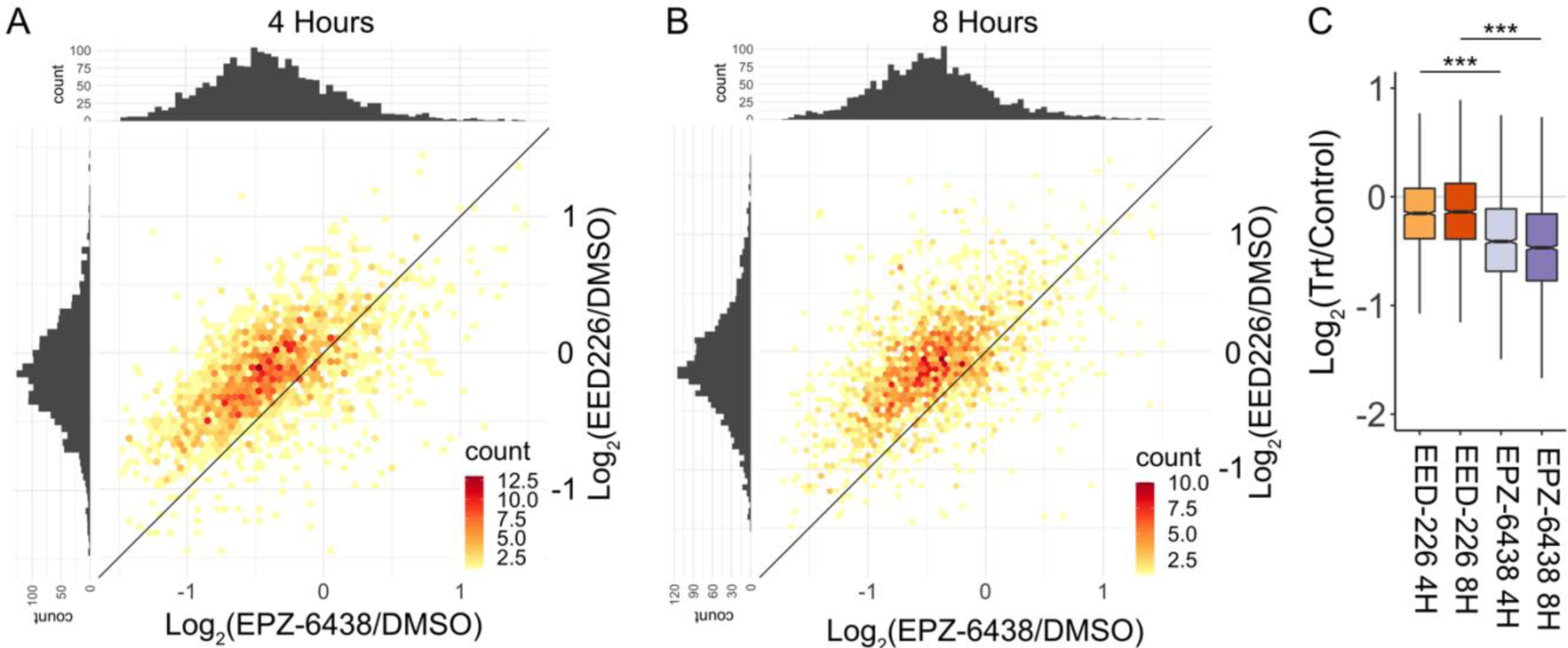
Differential effects at nucleation sites upon catalytic and allosteric inhibition of PRC2. **A**) Log2 ratio of the enrichment of H3K27me3 at nucleation sites comparing 4h treatment with PRC2 inhibitors to control (DMSO) is plotted as a heatmap with 2d hexagonal binning. The color scale mapping the number of sites in each bin is included in the bottom right corner of the plot. For each nucleation site, the gate with post-replication spike in each sample was used. **B**) Same as (**A**) for 8h treatment. **C**) Quantification of the H3K7me3 enrichment at nucleation sites at the post-replication spike from CUT&Flow, normalized to control (DMSO) is shown as a boxplot for each treatment. Significance string: ***p<2.2×10^−16^. P-values calculated using Wilcoxon rank sum test.

## Discussion

In this study, we have identified nucleation sites for every polycomb domain greater than 10 kb in size in mouse embryonic stem cells and show that rejuvenation of H3K27me3 every cell cycle occurs by nucleation and spreading. By analyzing a broad histone modification in the context of domains, we have shown that each H3K27me3 domain represents a microcosm of the genome-wide reader/writer/eraser kinetics. Using this microcosm model, we find polycomb domains feature a wide range of PRC2 kinetics. This range of PRC2 kinetics across domains has precluded previous attempts to identify nucleation sites genome-wide: any genome-wide criteria to identify nucleation sites would identify only the top enrichments and not segments in each domain that could nucleate.

Genetic studies and reporter assays have been traditionally used to identify developmental enhancers and PREs in *Drosophila*^16^. However, genetic deletions have proved notoriously difficult as a tool for identifying PREs in mammals presumably due to a high level of redundancy^17^. Here we show that kinetics of H3K27me3 in the timescale of hours can be a viable alternative to identify PRC2 nucleation sites both in the context of drug-washout and replication. We defined nucleation sites without relying on any steady-state chromatin properties, nevertheless our analyses tie together steady-state knowledge of Polycomb function with nucleation. These include enrichment of H3K27me3, Suz12, Ezh2, and accessory PRC2 subunits at nucleation sites compared to spreading sites, and significant overlap between nucleation sites and CpG islands and bivalent promoters.

We were able to uncover nucleation in drug washout studies because mESCs can both initiate and maintain most of their polycomb domains. A recent study has shown that differentiated cell types cannot regain some polycomb domains upon prolonged treatment with PRC2 inhibitor followed by drug washout^39^. Thus, using drug-washout kinetics is not a widely applicable method to identify nucleation sites across cell types. Here we show that CUT&Flow can be used to identify PRC2 nucleation in unperturbed cells. CUT&Flow is easily transferrable across cell types and can be used to ask if nucleation sites are preserved in polycomb domains that are retained upon differentiation of mESC. Importantly, CUT&Flow is able to capture the effect of altered PRC2 kinetics in short timescales. At just 4h of PRC2 inhibition, we can already observe the global slowdown in H3K27me3 restoration and loss of nucleation post-replication. The ability to capture fast changes in dynamics of H3K27me3 domains enabled us to distinguish the effects of catalytic inhibition compared to allosteric inhibition of PRC2 at nucleation sites, thus establishing new H3K27me3 methylation as the cause for the spike in H3K27me3 levels at nucleation sites post-replication. Finally, CUT&Flow also shows that the kinetics of *de novo* nucleation can explain the restoration of polycomb domains post-dilution due to replication.

Surprisingly, we find a subset of domains whose nucleation appears to happen pre-replication. Our findings mirror almost exactly what was observed for BX-C PRE in *Drosophila* cells, where nucleation occurred in the early S-phase before replication of the PRE in the late S-phase^40^. Our observations lead us to propose that the generation and maintenance of polycomb domains genome-wide depends on the locus-specific binding affinity and on-rates of PRC2 balanced by the replication dilution of parental histones. Such a model places reliance on starting states during differentiation and ongoing kinetic processes for setting polycomb domains in differentiated cells in addition to pure epigenetic memory. This is exemplified by recent observations that a high burst of transcription can erase polycomb memory in differentiated cells^39^. Our observations in this study thus suggest that polycomb memory is a product of constant nucleation in addition to the inheritance of parental histone modifications.

## Author Contributions

S.R. conceptualized this study. G.M.B.V. developed CUT&Flow method and performed the experiments. S.R. performed the data analysis. S.R. wrote the original draft of the manuscript. G.M.B.V. reviewed and edited the manuscript.

## Acknowledgements

We thank Dr. Jay Hesselberth for suggesting the idea that led to CUT&Flow. We thank Dr. Daphne Avgousti, Dr. David Bentley, Dr. Jay Hesselberth, Dr. Aaron Johnson, Dr. Olivia Rissland, Dr. Sujatha Jagannathan, and the members of the Ramachandran lab for critical reading of the manuscript and their insightful comments. We thank Christine Childs, Lester Acosta, and the CU Cancer Center Flow Cytometry Shared Resource (NIH grant P30CA046934) for their valuable assistance with the flow cytometry and sorting procedures. This work was supported by the RNA Bioscience Initiative, the University of Colorado School of Medicine, and NIH grant R35GM133434 (S. Ramachandran).

## Materials

All materials used in the study are listed in **Table S1**.

## Methods

### Cell lines and culture conditions

Mouse E14 embryonic STEM cells (mESCs) were maintained in complete medium consisting of KnockOut DMEM medium (Gibco) supplemented with 20% ES-qualified fetal bovine serum (FBS, Gibco), 2 mM Glutamax (Gibco), 10 U/ml penicillin-streptomycin (Gibco), 1X non-essential amino acids (Gibco), 1X 2-mercaptoethanol (Gibco) and 1000 U/ml Leukemia Inhibitory Factor (LIF, Millipore-Sigma). mESCs were seeded in 0.1% gelatin-coated culture flasks to a final concentration of 0.4 × 10^6^ cells/ml in complete medium and kept in a 37°C humidified incubator with 5% CO_2_.

### Inhibitor treatments

mESCs were seeded in 0.1% gelatin-coated 6-well plates to a final concentration of 0.4 × 10^6^ cells/ml in complete medium and incubated in a 37°C humidified incubator with 5% CO_2_ for 12 h before treatment. Medium was then replaced by fresh complete medium containing EPZ-6438 (MedChem Express) or EED226 (MedChem Express) in a final concentration of 10µM or 0.1% DMSO for untreated control. Plates were then incubated for 4h or 8h in a 37°C humidified incubator with 5% CO_2_, harvested by dissociation with 0.25% trypsin-EDTA (Gibco), and immediately processed for CUT&Flow and Flow Cytometry analysis, as described below.

### CUT&Flow

CUT&Flow was adapted from CUT&Tag ^31^. About 2,000,000 – 3,000,000 untreated or inhibitor-treated mESCs were collected per sample, in low-retention tubes. Samples were processed by centrifugation at 600 x g for 3 min at RT, in a swinging bucket centrifuge. Cells were lysed and nuclei extracted in nuclear extraction buffer (20 mM HEPES pH 7.9, 10 mM KCl, 0.1% Triton X-100, 20% glycerol, 0.5 mM spermidine and 1x Roche, cOmplete™, Mini, EDTA-free Protease Inhibitor Cocktail CAT# 04693159001) for 10 min on ice, and then subdivided into multiple 500,000 nuclei reactions. Each reaction was incubated with primary antibody (H3K27me3 or IgG, 1:100) in antibody 150 buffer (2 mM EDTA in Digitonin 150 buffer) for 1h at RT, followed by incubation with secondary antibody (Anti-Rabbit, 1:100) in Digitonin 150 buffer (20 mM HEPES pH 7.5, 150 mM NaCl, 0.5 mM spermidine, 1x protease inhibitor cocktail and 0.01% digitonin) for 30 min at RT. Nuclei were next washed once in Digitonin 150 buffer and incubated with pAG-Tn5 (Epicypher Inc., 1:20) in Digitonin 300 buffer (20 mM HEPES pH 7.5, 300 mM NaCl, 0.5 mM spermidine, 1x protease inhibitor cocktail and 0.01% digitonin) for 1h at RT. After one wash in Digitonin 300 buffer, pAG-Tn5 was activated and tagmentation performed by incubation in tagmentation buffer (20 mM HEPES pH 7.5, 300 mM NaCl, 0.5 mM spermidine, 1x protease inhibitor cocktail and 10 mM MgCl_2_) for for 1h at 37°C. Tagmented nuclei were combined back into one tube per sample and incubated in TAPS buffer (10 mM TAPS pH 8.5 and 0.2 mM EDTA) for 5 min at RT to stop the tagmentation reaction. Samples were then resuspended in Krishan solution ^41^ (3.8 mM sodium citrate; 46 µg/ml propidium iodide (PI); 0.01% NP-40; 10 µg/ml RNase A) and kept at 4°C, protected from light, for up to three days, until sorting.

PI-stained nuclei were sorted based on DNA content in an Astrios EQ instrument (Beckman Coulter). Five fractions were defined: 0-G1, G2-M and three equal subdivisions of the S phase, classified as Early S, Mid S and Late S. A range of 950 to 7000 cells were collected between the gates. Right after sorting, samples were incubated in SDS release buffer (10 mM TAPS pH 8.5 and 0.1% SDS) for nuclei lysis and release of tagmented chromatin for 1h at 58°C. SDS was neutralized by addition of SDS quench buffer (0.67% Triton-X 100 in nuclease-free H2O) and PCR performed with NEBNext High-Fidelity 2x PCR Master Mix (NEB, M0541L) and barcoded primers for library construction, with the following protocol: 5 min at 58°C, 5 min at 72°C, 45 sec at 98°C, 18 cycles of 15 sec at 98°C and 10 sec at 60°C, followed by 1 min at 72°C for final extension.

### Flow Cytometry Analysis

500,000 untreated or inhibitor-treated mESCs were collected per sample, in low-retention tubes, and nuclei extraction was performed as described in the CUT&Flow section. Samples were incubated with primary antibody (H3K27me3 1:100) in antibody 150 buffer overnight at 4°C, followed by one wash with Digitonin 150 buffer and incubation with secondary antibody (CF640R, 1 drop) at RT for 30 min. Nuclei were next washed with Digitonin 150 buffer, resuspended in Krishan solution (Krishan, 1975), and kept at 4°C, protected from light, overnight, until analysis. Stained samples were run in Gallios 561 flow cytometer (Beckman Coulter) using 488 nm laser for PI and 638 nm laser for CF640R. The obtained fcs files were analyzed using FlowJo and Rstudio softwares.

### Analysis

All publicly available datasets used in this study are listed in **Table S2**.

### Defining Polycomb domains in mESCs

For Figure 1, published datasets from ^14^ were aligned to mm10 version of the *Mus musculus* genome. Coverage at 25 bp windows genome-wide was calculated as number of reads that mapped at that window, normalized by the factor N:

> N = 2800000000/(Total number of mapped reads)

2800000000 was a number chosen arbitrarily in the range of total mapped base pairs in M. musculus genome. The normalized read density in 25 bp bins were then smoothed with a running average spanning +/− 1000 bp around each bin. Domains were called by linking adjacent windows that satisfied two cutoffs: 4-fold enrichment over input and 2-fold enrichment over the genome-wide median. To account for short disruptions due to mappability issues, jumps of up to 750 bp were allowed while linking windows.

### Determining H3K27me3 recovery kinetics from drug washout datasets

Control H3K27me3 ChIP-seq dataset from^8^ was used for domain calling as described above. Then domains that overlapped between Ref.^8^ and Ref.^14^ datasets were used for all further analyses. The 7-day inhibition dataset, and datasets following washout for the following time points were used: 4, 8, 16, 24, 48, 96 hours^8^. Kinetics was analyzed at each 25 bp window of a domain. The H3K27me3 enrichment was first normalized at each window by the enrichment at 96h time point for all time points, so that the 96h time point would be 1 across all windows. After normalizing to final time-point, the time series matrix for each domain (an example shown in **Figure 2A**) was subjected to k-means clustering with k=7 in R^42^. The average time profile of H3K27me3 enrichment was calculated for each kinetic cluster and the time to reach 40% of recovery relative to 96-hour time point was interpolated for each kinetic cluster. This time, T_40_ was used to order the kinetic clusters from slowest to fastest. The sites corresponding to fastest (cluster 7) were used as nucleation sites for that domain for all further analyses.

### CUT&Flow analysis

CUT&Flow datasets were aligned to the mm10 version of *Mus musculus* genome using bowtie2, with the following parameters:

> --local --very-sensitive-local --no-unal --no-mixed --no-discordant -I 10 -X 2000

Samtools^43^ and bedtools^44^ were used for processing aligned reads from sam to bed files. Duplicate reads were discarded for further analysis if the reads had same start and end coordinates. Coverage at 100 bp windows genome-wide was calculated as number of reads that mapped at that window, normalized by the factor N:

> N = 2800000000/(Total number of mapped reads)

2800000000 was a number chosen arbitrarily in the range of total mapped base pairs in M. musculus genome. For replication timing, the final data matrix for mESC from Ref.^32^ was used. The data matrix contains genomic intervals and for each interval, the percentage of replication in 16 S-phase windows. The S-phase window with highest percentage of replication was used for the plots shown in Figure 3E and Figure 4C. For Figure 2D, the average normalized read count from the whole domain for all gates were calculated. The gate with lowest normalized read count that was lower by at least 0.5 than the next lowest gate was assigned to the domain. For the heatmap, the values for all gates of a domain were normalized by the highest value and log2 transformed.

For Figure 4A, the average normalized read count at each nucleation site was first divided by the average normalized read count at its corresponding spreading sites to get the nucleation site enrichment ratio for each gate. The gate with highest the nucleation site enrichment ratio that was higher by at least 0.5 than the ratio at the lowest gate and by 0.2 than the ratio at the next highest gate was assigned to the nucleation site. If a gate satisfying the above criteria was not found, it went to the unassigned category. For the heatmap, row-wise Z-score of the nucleation site enrichment ratio was plotted for each nucleation site, where the nucleation sites were ordered by the assigned gates. For Figures 5D-F, we assigned replication timing for domains as follows: Early S if %replication peaked in S-phase window <6, Mid S if replication peaked in S-phase window between 6 and 10, and late S if replication peaked in S-phase window > 10. The gene ontology overrepresentation analysis was performed using the WebGestalt server^45^.

## References

1. Francis, N.J., and Kingston, R.E. (2001). Mechanisms of transcriptional memory. Nat Rev Mol Cell Biol 2, 409–421. 10.1038/35073039.

2. Bowman, S.K., Deaton, A.M., Domingues, H., Wang, P.I., Sadreyev, R.I., Kingston, R.E., and Bender, W. (2014). H3K27 modifications define segmental regulatory domains in the Drosophila bithorax complex. Elife 3, e02833. 10.7554/eLife.02833.

3. Kundu, S., Ji, F., Sunwoo, H., Jain, G., Lee, J.T., Sadreyev, R.I., Dekker, J., and Kingston, R.E. (2017). Polycomb Repressive Complex 1 Generates Discrete Compacted Domains that Change during Differentiation. Mol Cell 65, 432–446 e435. 10.1016/j.molcel.2017.01.009.

4. Cao, R., Wang, L., Wang, H., Xia, L., Erdjument-Bromage, H., Tempst, P., Jones, R.S., and Zhang, Y. (2002). Role of histone H3 lysine 27 methylation in Polycomb-group silencing. Science 298, 1039–1043. 10.1126/science.1076997.

5. Kuzmichev, A., Nishioka, K., Erdjument-Bromage, H., Tempst, P., and Reinberg, D. (2002). Histone methyltransferase activity associated with a human multiprotein complex containing the Enhancer of Zeste protein. Genes Dev 16, 2893–2905. 10.1101/gad.1035902.

6. Muller, J., Hart, C.M., Francis, N.J., Vargas, M.L., Sengupta, A., Wild, B., Miller, E.L., O’Connor, M.B., Kingston, R.E., and Simon, J.A. (2002). Histone methyltransferase activity of a Drosophila Polycomb group repressor complex. Cell 111, 197–208. 10.1016/s0092-8674(02)00976-5.

7. Hojfeldt, J.W., Hedehus, L., Laugesen, A., Tatar, T., Wiehle, L., and Helin, K. (2019). Non-core Subunits of the PRC2 Complex Are Collectively Required for Its Target-Site Specificity. Mol Cell 76, 423–436 e423. 10.1016/j.molcel.2019.07.031.

8. Hojfeldt, J.W., Laugesen, A., Willumsen, B.M., Damhofer, H., Hedehus, L., Tvardovskiy, A., Mohammad, F., Jensen, O.N., and Helin, K. (2018). Accurate H3K27 methylation can be established de novo by SUZ12-directed PRC2. Nat Struct Mol Biol 25, 225–232. 10.1038/s41594-018-0036-6.

9. Margueron, R., Justin, N., Ohno, K., Sharpe, M.L., Son, J., Drury, W.J., 3rd, Voigt, P., Martin, S.R., Taylor, W.R., De Marco, V., et al. (2009). Role of the polycomb protein EED in the propagation of repressive histone marks. Nature 461, 762–767. 10.1038/nature08398.

10. Shao, Z., Raible, F., Mollaaghababa, R., Guyon, J.R., Wu, C.T., Bender, W., and Kingston, R.E. (1999). Stabilization of chromatin structure by PRC1, a Polycomb complex. Cell 98, 37–46. 10.1016/S0092-8674(00)80604-2.

11. Wang, H., Wang, L., Erdjument-Bromage, H., Vidal, M., Tempst, P., Jones, R.S., and Zhang, Y. (2004). Role of histone H2A ubiquitination in Polycomb silencing. Nature 431, 873–878. 10.1038/nature02985.

12. Blackledge, N.P., Fursova, N.A., Kelley, J.R., Huseyin, M.K., Feldmann, A., and Klose, R.J. (2020). PRC1 Catalytic Activity Is Central to Polycomb System Function. Mol Cell 77, 857–874 e859. 10.1016/j.molcel.2019.12.001.

13. Fursova, N.A., Blackledge, N.P., Nakayama, M., Ito, S., Koseki, Y., Farcas, A.M., King, H.W., Koseki, H., and Klose, R.J. (2019). Synergy between Variant PRC1 Complexes Defines Polycomb-Mediated Gene Repression. Mol Cell 74, 1020–1036 e1028. 10.1016/j.molcel.2019.03.024.

14. Banaszynski, L.A., Wen, D., Dewell, S., Whitcomb, S.J., Lin, M., Diaz, N., Elsasser, S.J., Chapgier, A., Goldberg, A.D., Canaani, E., et al. (2013). Hira-dependent histone H3.3 deposition facilitates PRC2 recruitment at developmental loci in ES cells. Cell 155, 107–120. 10.1016/j.cell.2013.08.061.

15. Deal, R.B., Henikoff, J.G., and Henikoff, S. (2010). Genome-wide kinetics of nucleosome turnover determined by metabolic labeling of histones. Science 328, 1161–1164. 10.1126/science.1186777.

16. Steffen, P.A., and Ringrose, L. (2014). What are memories made of? How Polycomb and Trithorax proteins mediate epigenetic memory. Nat Rev Mol Cell Biol 15, 340–356. 10.1038/nrm3789.

17. Schorderet, P., Lonfat, N., Darbellay, F., Tschopp, P., Gitto, S., Soshnikova, N., and Duboule, D. (2013). A genetic approach to the recruitment of PRC2 at the HoxD locus. PLoS Genet 9, e1003951. 10.1371/journal.pgen.1003951.

18. Woo, C.J., Kharchenko, P.V., Daheron, L., Park, P.J., and Kingston, R.E. (2010). A region of the human HOXD cluster that confers polycomb-group responsiveness. Cell 140, 99–110. 10.1016/j.cell.2009.12.022.

19. Sing, A., Pannell, D., Karaiskakis, A., Sturgeon, K., Djabali, M., Ellis, J., Lipshitz, H.D., and Cordes, S.P. (2009). A vertebrate Polycomb response element governs segmentation of the posterior hindbrain. Cell 138, 885–897. 10.1016/j.cell.2009.08.020.

20. Simon, J., Chiang, A., Bender, W., Shimell, M.J., and O’Connor, M. (1993). Elements of the Drosophila bithorax complex that mediate repression by Polycomb group products. Dev Biol 158, 131–144. 10.1006/dbio.1993.1174.

21. Ringrose, L., and Paro, R. (2004). Epigenetic regulation of cellular memory by the Polycomb and Trithorax group proteins. Annu Rev Genet 38, 413–443. 10.1146/annurev.genet.38.072902.091907.

22. Oksuz, O., Narendra, V., Lee, C.H., Descostes, N., LeRoy, G., Raviram, R., Blumenberg, L., Karch, K., Rocha, P.P., Garcia, B.A., et al. (2018). Capturing the Onset of PRC2-Mediated Repressive Domain Formation. Mol Cell 70, 1149–1162 e1145. 10.1016/j.molcel.2018.05.023.

23. Kuroda, M.I., Kang, H., De, S., and Kassis, J.A. (2020). Dynamic Competition of Polycomb and Trithorax in Transcriptional Programming. Annu Rev Biochem 89, 235–253. 10.1146/annurev-biochem-120219-103641.

24. Alabert, C., Barth, T.K., Reveron-Gomez, N., Sidoli, S., Schmidt, A., Jensen, O.N., Imhof, A., and Groth, A. (2015). Two distinct modes for propagation of histone PTMs across the cell cycle. Genes Dev 29, 585–590. 10.1101/gad.256354.114.

25. Escobar, T.M., Oksuz, O., Saldana-Meyer, R., Descostes, N., Bonasio, R., and Reinberg, D. (2019). Active and Repressed Chromatin Domains Exhibit Distinct Nucleosome Segregation during DNA Replication. Cell 179, 953–963 e911. 10.1016/j.cell.2019.10.009.

26. Gaydos, L.J., Wang, W., and Strome, S. (2014). Gene repression. H3K27me and PRC2 transmit a memory of repression across generations and during development. Science 345, 1515–1518. 10.1126/science.1255023.

27. Reveron-Gomez, N., Gonzalez-Aguilera, C., Stewart-Morgan, K.R., Petryk, N., Flury, V., Graziano, S., Johansen, J.V., Jakobsen, J.S., Alabert, C., and Groth, A. (2018). Accurate Recycling of Parental Histones Reproduces the Histone Modification Landscape during DNA Replication. Mol Cell 72, 239–249 e235. 10.1016/j.molcel.2018.08.010.

28. Wang, X., Paucek, R.D., Gooding, A.R., Brown, Z.Z., Ge, E.J., Muir, T.W., and Cech, T.R. (2017). Molecular analysis of PRC2 recruitment to DNA in chromatin and its inhibition by RNA. Nat Struct Mol Biol 24, 1028–1038. 10.1038/nsmb.3487.

29. Yuan, W., Wu, T., Fu, H., Dai, C., Wu, H., Liu, N., Li, X., Xu, M., Zhang, Z., Niu, T., et al. (2012). Dense chromatin activates Polycomb repressive complex 2 to regulate H3 lysine 27 methylation. Science 337, 971–975. 10.1126/science.1225237.

30. Wang, Y., Long, H., Yu, J., Dong, L., Wassef, M., Zhuo, B., Li, X., Zhao, J., Wang, M., Liu, C., et al. (2018). Histone variants H2A.Z and H3.3 coordinately regulate PRC2-dependent H3K27me3 deposition and gene expression regulation in mES cells. BMC Biol 16, 107 10.1186/s12915-018-0568-6.

31. Kaya-Okur, H.S., Wu, S.J., Codomo, C.A., Pledger, E.S., Bryson, T.D., Henikoff, J.G., Ahmad, K., and Henikoff, S. (2019). CUT&Tag for efficient epigenomic profiling of small samples and single cells. Nat Commun 10, 1930 10.1038/s41467-019-09982-5.

32. Zhao, P.A., Sasaki, T., and Gilbert, D.M. (2020). High-resolution Repli-Seq defines the temporal choreography of initiation, elongation and termination of replication in mammalian cells. Genome Biology 21, 76 10.1186/s13059-020-01983-8.

33. Flury, V., Reveron-Gomez, N., Alcaraz, N., Stewart-Morgan, K.R., Wenger, A., Klose, R.J., and Groth, A. (2023). Recycling of modified H2A-H2B provides short-term memory of chromatin states. Cell 186, 1050–1065 e1019. 10.1016/j.cell.2023.01.007.

34. Asenjo, H.G., Gallardo, A., Lopez-Onieva, L., Tejada, I., Martorell-Marugan, J., Carmona-Saez, P., and Landeira, D. (2020). Polycomb regulation is coupled to cell cycle transition in pluripotent stem cells. Sci Adv 6, eaay4768. 10.1126/sciadv.aay4768.

35. Mendenhall, E.M., Koche, R.P., Truong, T., Zhou, V.W., Issac, B., Chi, A.S., Ku, M., and Bernstein, B.E. (2010). GC-rich sequence elements recruit PRC2 in mammalian ES cells. PLOS Genetics 6, e1001244. 10.1371/journal.pgen.1001244.

36. Riising, E.M., Comet, I., Leblanc, B., Wu, X., Johansen, J.V., and Helin, K. (2014). Gene Silencing Triggers Polycomb Repressive Complex 2 Recruitment to CpG Islands Genome Wide. Molecular Cell 55, 347–360. 10.1016/j.molcel.2014.06.005.

37. Knutson, S.K., Kawano, S., Minoshima, Y., Warholic, N.M., Huang, K.C., Xiao, Y., Kadowaki, T., Uesugi, M., Kuznetsov, G., Kumar, N., et al. (2014). Selective inhibition of EZH2 by EPZ-6438 leads to potent antitumor activity in EZH2-mutant non-Hodgkin lymphoma. Mol Cancer Ther 13, 842–854. 10.1158/1535-7163.MCT-13-0773.

38. Qi, W., Zhao, K., Gu, J., Huang, Y., Wang, Y., Zhang, H., Zhang, M., Zhang, J., Yu, Z., Li, L., et al. (2017). An allosteric PRC2 inhibitor targeting the H3K27me3 binding pocket of EED. Nat Chem Biol 13, 381–388. 10.1038/nchembio.2304.

39. Holoch, D., Wassef, M., Lovkvist, C., Zielinski, D., Aflaki, S., Lombard, B., Hery, T., Loew, D., Howard, M., and Margueron, R. (2021). A cis-acting mechanism mediates transcriptional memory at Polycomb target genes in mammals. Nat Genet 53, 1686–1697. 10.1038/s41588-021-00964-2.

40. Lanzuolo, C., Lo Sardo, F., Diamantini, A., and Orlando, V. (2011). PcG complexes set the stage for epigenetic inheritance of gene silencing in early S phase before replication. PLoS Genet 7, e1002370. 10.1371/journal.pgen.1002370.

41. Krishan, A. (1975). Rapid flow cytofluorometric analysis of mammalian cell cycle by propidium iodide staining. J Cell Biol 66, 188–193. 10.1083/jcb.66.1.188.

42. RCoreTeam (2019). R: A Language and Environment for Statistical Computing (R Foundation for Statistical Computing).

43. Li, H., Handsaker, B., Wysoker, A., Fennell, T., Ruan, J., Homer, N., Marth, G., Abecasis, G., Durbin, R., and Genome Project Data Processing, S. (2009). The Sequence Alignment/Map format and SAMtools. Bioinformatics 25, 2078–2079. 10.1093/bioinformatics/btp352.

44. Quinlan, A.R., and Hall, I.M. (2010). BEDTools: a flexible suite of utilities for comparing genomic features. Bioinformatics 26, 841–842. 10.1093/bioinformatics/btq033.

45. Liao, Y., Wang, J., Jaehnig, E.J., Shi, Z., and Zhang, B. (2019). WebGestalt 2019: gene set analysis toolkit with revamped UIs and APIs. Nucleic Acids Res 47, W199–W205. 10.1093/nar/gkz401.

