## Supplementary Material for "Nucleation and spreading rejuvenate polycomb domains every cell cycle"

### Supplemental Materials

**Figure S1**

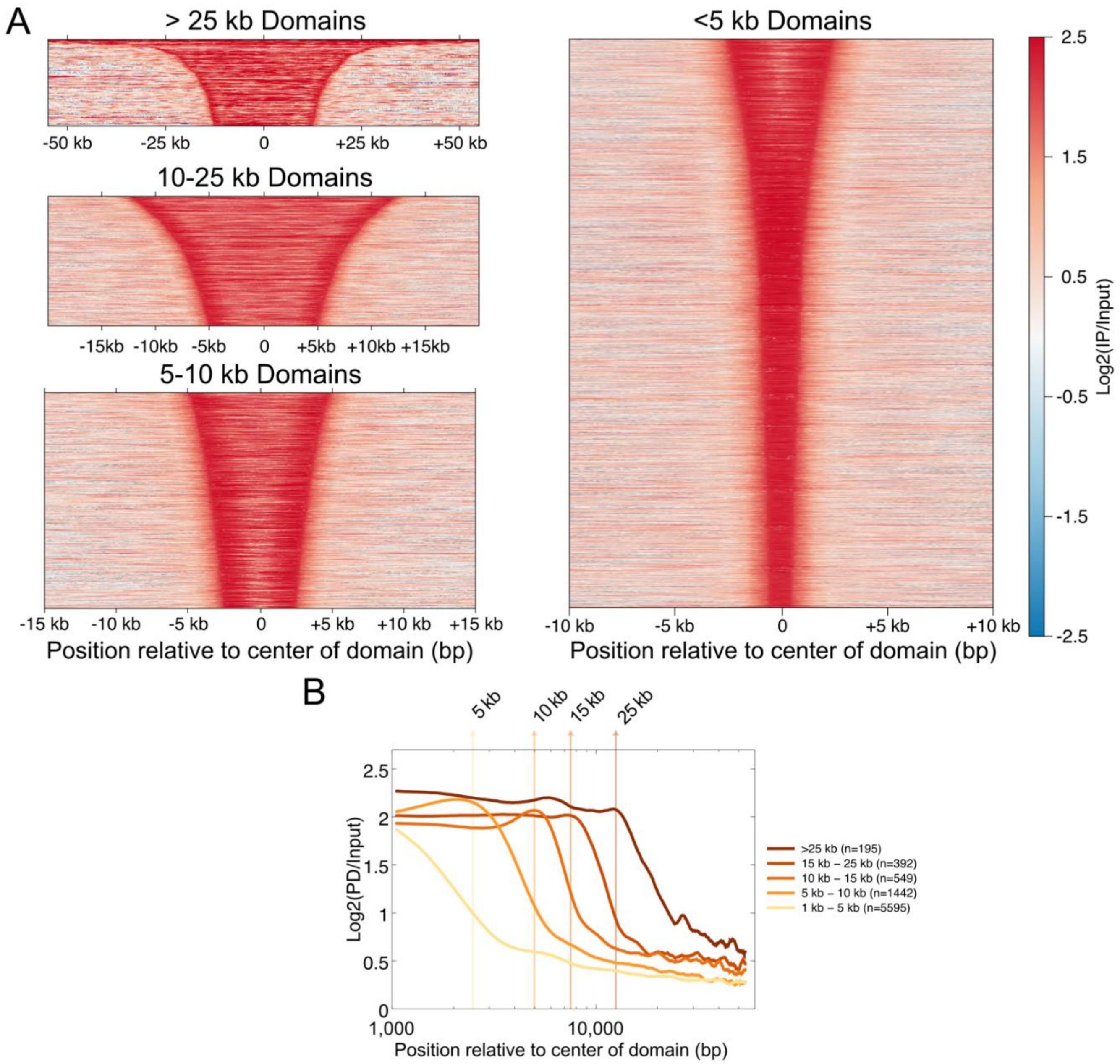


**Figure S1. Genome-wide identification of H3K27me3 domains. A**) Heatmaps of H3K27me3 ChIP enrichment over input plotted for domains relative to their center, at various scales. **B**) Average profile of H3K27me3 ChIP enrichment over input in the log scale plotted for domains relative to their center, to demonstrate domain edges. The position relative to domain center is plotted on a log scale.

**Figure S2**

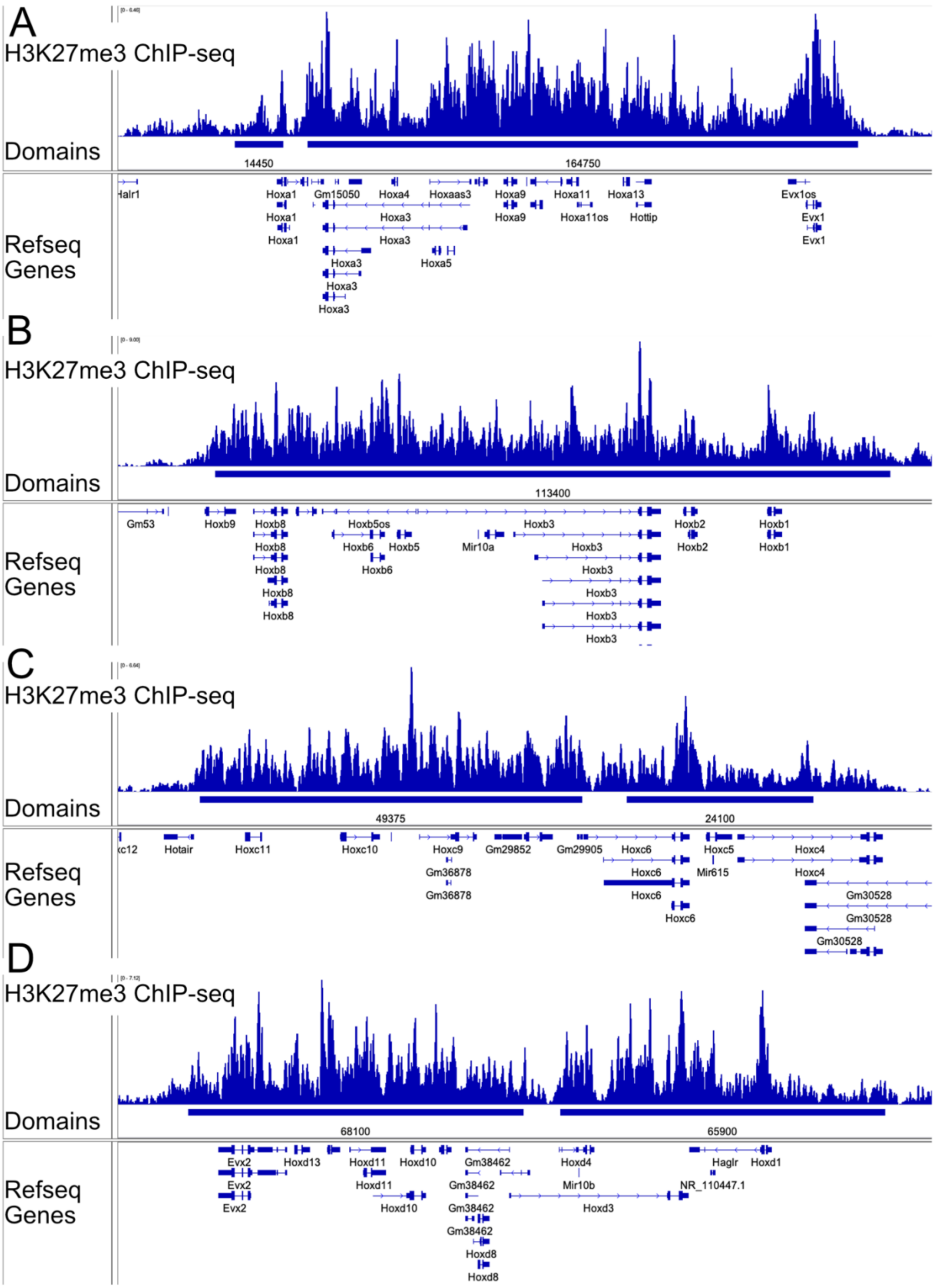

A
H3K27me3 ChIP-seq
Domains
14450
164750
Refseq Genes
Halr1
Hoxa1
Gm15050
Hoxa4
Hoxaas3
Hoxa9
Hoxa11
Hoxa13
Evx1os
Hoxa1
Hoxa3
Hoxa9
Hoxa11os
Hottip
Evx1
Hoxa1
Hoxa3
Evx1
Hoxa3
Hoxa5
Hoxa3
Hoxa3
B
H3K27me3 ChIP-seq
Domains
113400
Refseq Genes
Gm53
Hoxb9
Hoxb8
Hoxb5os
Hoxb3
Hoxb2
Hoxb1
Hoxb8
Hoxb6
Hoxb5
Mir10a
Hoxb3
Hoxb2
Hoxb1
Hoxb8
Hoxb6
Hoxb3
Hoxb8
Hoxb3
Hoxb8
Hoxb3
Hoxb3
C
H3K27me3 ChIP-seq
Domains
49375
24100
Refseq Genes
Hotair
Hoxc11
Hoxc10
Hoxc9
Gm29852
Gm29905
Hoxc6
Hoxc5
Hoxc4
Gm36878
Hoxc6
Mir615
Hoxc4
Gm36878
Hoxc6
Gm30528
Hoxc6
Gm30528
Gm30528
D
H3K27me3 ChIP-seq
Domains
68100
65900
Refseq Genes
Evx2
Hoxd13
Hoxd11
Hoxd10
Gm38462
Hoxd4
Hagir
Hoxd1
Evx2
Hoxd11
Gm38462
Mir10b
NR_110447.1
Evx2
Hoxd10
Gm38462
Hoxd3
Gm38462
Hoxd8
Hoxd8

**Figure S2. Large domains defined at Hox loci. A**) Our algorithm identifies 164 kb domain at the Hoxa locus (chr6:52,120,00-52,340,000). **B**) A 113 kb domain is defined at the Hoxb locus (chr11:96,260,000-96,390). **C**) Two domains of 49 kb and 24 kb respectively are defined at the HoxC locus (chr15:102,940,000-103,040,000). **D**) Two domains of 68 kb and 66 kb respectively are defined at the HoxD locus (chr2:74,640,000-74,790,000).

**Figure S3**

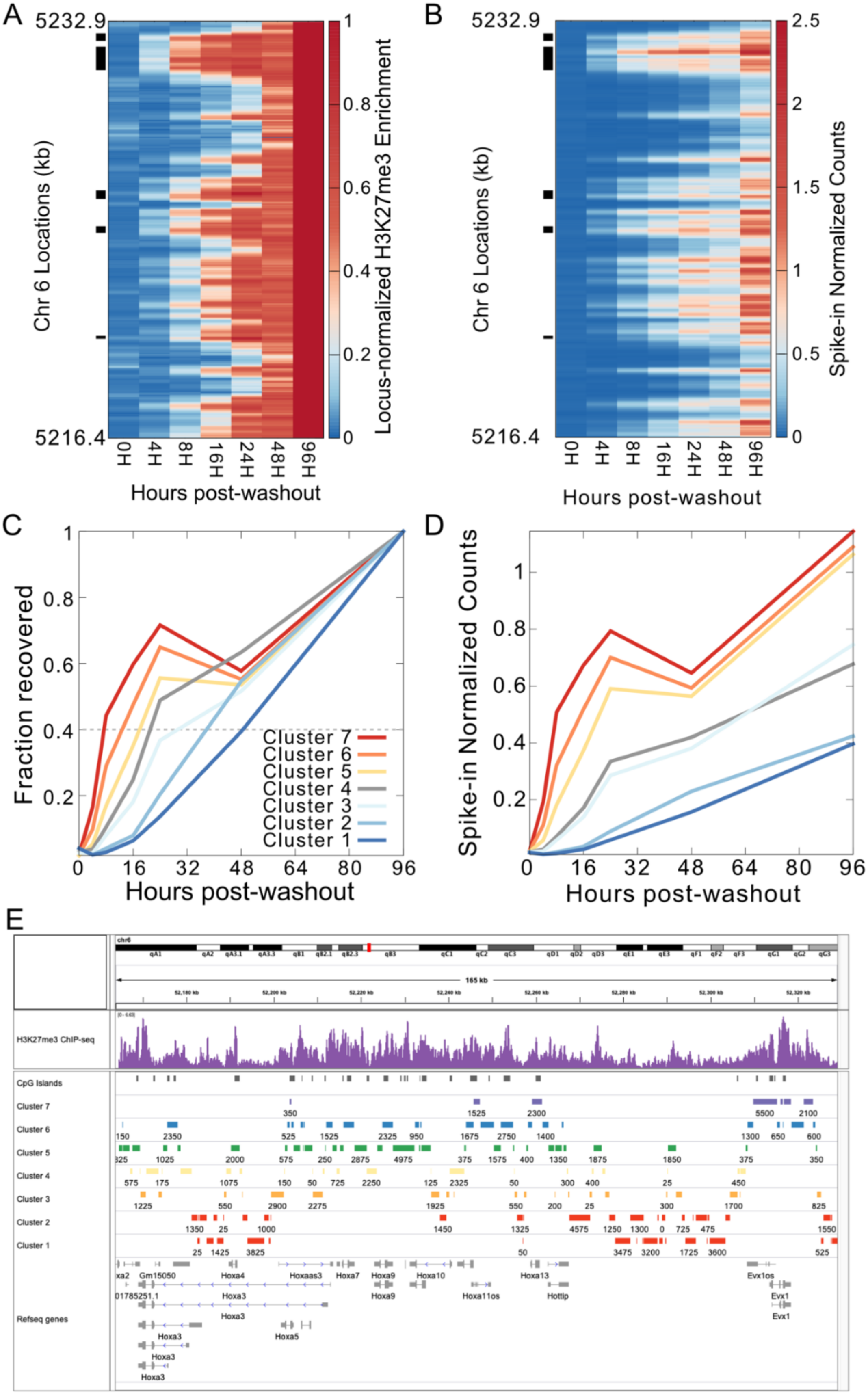

A
B
C
D
E
5232.9
5216.4
Chr 6 Locations (kb)
0H
4H
8H
16H
24H
48H
96H
Hours post-washout
Locus-normalized H3K27me3 Enrichment
Spike-in Normalized Counts
Fraction recovered
Cluster 7
Cluster 6
Cluster 5
Cluster 4
Cluster 3
Cluster 2
Cluster 1
chr6
165 kb
H3K27me3 ChIP-seq
CpG Islands
Refseq genes
Gm15050
Hoxa4
Hoxaas3
Hoxa7
Hoxa9
Hoxa10
Hoxa13
Evx1os
Hoxa3
Hoxa11os
Hottip
Evx1
Hoxa5

**Figure S3. Intradomain kinetics of H3K27me3 recovery at the Hoxa domain. A)** Heatmap of H3K27me3 ChIP enrichment at time points post-washout of Ezh2 inhibitor (x axis) normalized to the ChIP enrichment at 96 hours post-washout. Each horizontal line of the heatmap represents 25 bp step of the genomic location on the Hoxa domain. **B**) Same as (**A**) but not normalized to the ChIP enrichment at 96 hours post-washout. The black bars to the left of the plots in (**A**) and (**B**) are the identified nucleation sites. **C**) Average H3K27me3 ChIP enrichment of each of the 7 clusters obtained by k-means clustering as a function of time post-washout, normalized to the ChIP enrichment at 96 hours post-washout. **D**) Average H3K27me3 ChIP enrichment of each of the 7 clusters obtained by k-means clustering as a function of time post-washout, not normalized to the ChIP enrichment at 96 hours post-washout. **E**) Genome browser snapshot showing enrichment of H3K27me3 ChIP-seq, CpG islands, the kinetic clusters 1-7, where cluster 7 represents nucleation sites, and Refseq genes.

**Figure S4**

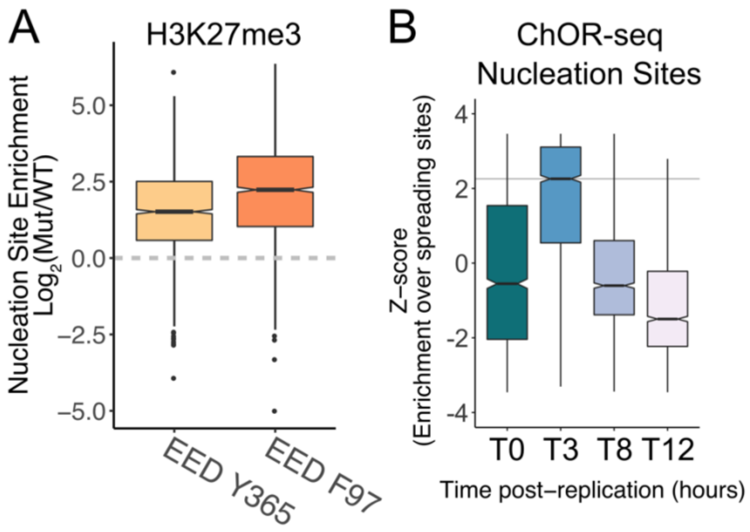


**Figure S4. EED cage mutants and ChOR-seq show increased enrichment of H3K27me3 at nucleation sites. A**) Nucleation strength, the change in enrichment of H3K27me3 at nucleation sites relative to spreading sites in EED cage mutants (data from Oksuz *et al.*), shown as a boxplot across domains. **B**) Distribution of the Z-score of nucleation site enrichment over spreading sites at different times post-replication shown as a boxplot (using ChOR-seq data from Flury *et al.*). **C**) The enrichment of H3K27me3 at clusters 3-7 over the enrichment at spreading sites (clusters 1,2) plotted as Z-scores.

**Figure S5**

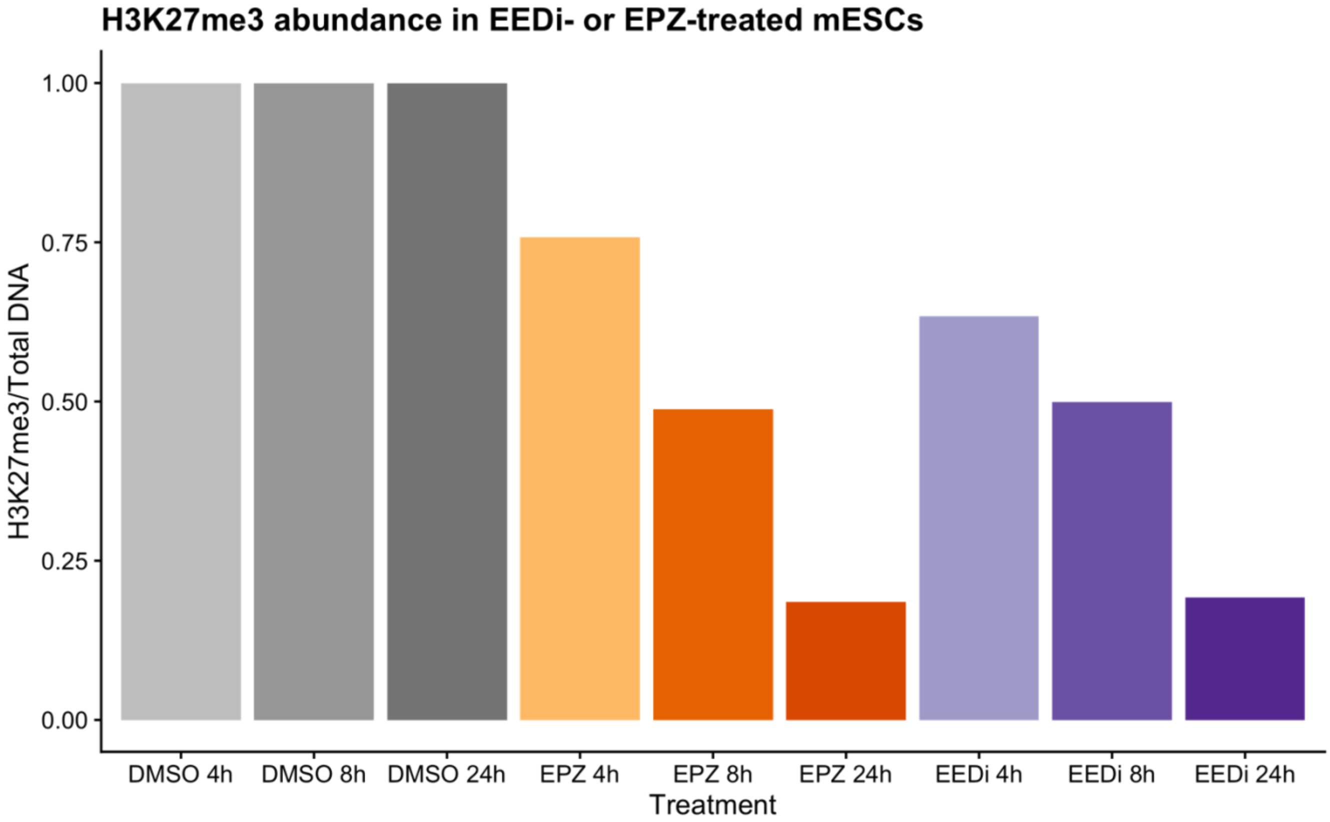


**Figure S5. Decrease in global H3K27me3 levels upon inhibiting PRC2.** Median H3K27me3 levels normalized to control (DMSO) samples plotted for various treatment times. Total H3K27me3 and DNA levels were determined by flow cytometry.

### Table S1. Critical resources used in this study

| Resource | Reference |
|---|---|
| **Cell lines** | |
| E14 mouse embryonic stem cells | Dr. Peter J. Koch |
| **Antibodies** | |
| Anti-trimethyl-Histone H3 (Lys27) C36B11 rabbit monoclonal (0.1 mg/ml) | Cell Signaling Technology; Cat # 9733; RRID: AB_2616029 |
| Normal IgG rabbit polyclonal (1 mg/ml) | R&D Biosciences; Cat # AB-105-C; RRID: AB_354266 |
| Anti-rabbit goat mixed monoclonal (1 mg/ml) | Epicypher; Cat # 13-0047 |
| CF640R donkey anti-rabbit IgG | Biotium; Cat # 20962 |
| **Chemicals** | |
| Tazemetostat/ EPZ-6438 | MedChemExpress; Cat # HY-13803 |
| EED226 | MedChemExpress; Cat # HY-101117 |
| 2% gelatin solution | Millipore-Sigma; Cat # G1393 |
| KnockOut DMEM | Gibco; Cat # 10829018 |
| FBS, ES-qualified | Gibco; Cat # 16141079 |
| Glutamax (100X) | Gibco; Cat # 35050061 |
| Penicillin-streptomycin (10,000 U/ml) | Gibco; Cat # 15140122 |
| MEM non-essential amino acids (100X) | Gibco; Cat # 11140050 |
| 2-mercaptoethanol (1000X) | Gibco; Cat # 21985023 |
| ESGRO-LIF | Millipore-Sigma; Cat # ESG1107 |
| Trypsin-EDTA (0.25%), phenol red | Gibco; Cat# 25200056 |
| Digitonin | Millipore-Sigma; Cat # 300410 |
| Spermidine | Millipore-Sigma; Cat # S2626 |
| Roche Complete Protease Inhibitor mini EDTA-Free tablets | Millipore-Sigma; Cat # 04693159001 |
| TAPS buffer 0.2 M | Boston Scientific; Cat # BB-2372 |
| CUTANA pAG-Tn5 | Epicypher; Cat # 15-1017 |
| AMPure XP SPRI beads | Beckman Coulter; Cat # A63881 |
| Propidium Iodide | Millipore-Sigma; Cat # P4170 |
| RNase A, DNase and protease-free (10 mg/mL) | ThermoFisher; Cat # EN0531 |
| **Commercial Assays** | |
| NEBNext High-Fidelity 2× PCR Master Mix | NEB; Cat # M0541 |
| Qubit dsDNA HS Assay Kit | ThermoFisher; Cat # Q32854 |
| Tapestation High Sensitivity D1000 ScreenTapes | Agilent; Cat # 5067-5584 |
| Tapestation High Sensitivity D1000 Sample Buffer & Ladder | Agilent; Cat # 5067-5585 |
| **Oligonucleotides** | |
| Universal i5 primer for CUT&tag: 5’ AATGATACGGCGACCACCGAGATCTACACTAGATCGCTCGTCGGCAGCGTCAGATGTGTAT 3’ | Epicypher |
| Barcoded i7 primers for CUT&Tag: 5' CAAGCAGAAGACGGCATACGAGATNNNNNNNNGTCTCGTGGGCTCGGAGATGTG 3' | Epicypher |
| **Equipments** | |

| | |
|---|---|
| Countess 3 Automated Cell Counter | ThermoFisher; Cat # AMQAX2000 |
| Magnetic Separation Rack, 0.2 mL Tubes | Epicypher; Cat # 10-0008 |
| 4200 TapeStation System | Agilent; Cat # G2991BA |
| Qubit 4 Fluorometer | ThermoFisher; Cat # Q33238 |
| Gallios 561 Flow Cytometer | Beckman Coulter |
| MoFlo Astrios EQ Cell Sorter | Beckman Coulter; Cat # B52102 |

**Table S2. External datasets used in this study, Related to STAR Methods**

| S. No. | Dataset | Figures | GEO Accession ID | Reference |
|---|---|---|---|---|
| 1 | H3K27me3 ChIPseq | Figure S1, S2 | GSM1033638 | [1] |
| 2 | H3K27me3 ChIPseq | Figure S1, S2 | GSM487549 | [1] |
| 3 | H3K27me3 ChIPseq | Figure 1 | GSM2779214 | [2] |
| 4 | H3K27me3 ChIPseq, 7-day treatment with EPZ6438 | Figure 1 | GSM2779215 | [2] |
| 5 | H3K27me3 ChIPseq 4 hours after washout of EPZ6438 | Figure 1 | GSM2779216 | [2] |
| 6 | H3K27me3 ChIPseq 8 hours after washout of EPZ6438 | Figure 1 | GSM2779217 | [2] |
| 7 | H3K27me3 ChIPseq 16 hours after washout of EPZ6438 | Figure 1 | GSM2779218 | [2] |
| 8 | H3K27me3 ChIPseq 24 hours after washout of EPZ6438 | Figure 1 | GSM2779219 | [2] |
| 9 | H3K27me3 ChIPseq 48 hours after washout of EPZ6438 | Figure 1 | GSM2779220 | [2] |
| 10 | H3K27me3 ChIPseq 96 hours after washout of EPZ6438 | Figure 1 | GSM2779221 | [2] |
| 11 | Jarid2 ChIP-seq | Figure 5 | GSM3021208 | [3] |
| 12 | Jarid2 ChIP-seq | Figure 5 | GSM3021209 | [3] |
| 13 | PCL2 ChIP-seq | Figure 5 | GSM2472747 | [3] |
| 14 | PCL2 ChIP-seq | Figure 5 | GSM2472748 | [3] |
| 15 | Ezh2 ChIP-seq | Figure 5 | GSM2472741 | [3] |
| 16 | Ezh2 ChIP-seq | Figure 5 | GSM2472742 | [3] |
| 17 | Suz12 ChIP-seq | Figure 5 | GSM2779242 | [2] |
| 18 | H2AK119ubiquitin ChIP-seq | Figure 5 | GSE119618 | [4] |
| 19 | ATAC-seq | Figure 5 | GSM1941479 | [5] |
| 20 | ATAC-seq | Figure 5 | GSM1941480 | [5] |
| 21 | ATAC-seq | Figure 5 | GSM1941487 | [5] |
| 22 | ATAC-seq | Figure 5 | GSM1941488 | [5] |
| 23 | H3.3 ChIP-seq | Figure 5 | GSM423355 | [6] |
| 24 | H3.3 ChIP-seq | Figure 5 | GSM1207786 | [1] |
| 25 | H3.3 ChIP-seq | Figure 5 | GSM487544 | [6] |
| 26 | H3.3 ChIP-seq | Figure 5 | GSM487551 | [6] |
| 27 | H3K27me3 ChIP-seq (EED Y365A) | Figure S4 | GSM2475241 | [7] |
| 28 | H3K27me3 ChIP-seq (EED Y365A) | Figure S4 | GSM2475248 | [7] |
| 29 | H3K27me3 ChIP-seq (EED F97A) | Figure S4 | GSM2475240 | [7] |
| 30 | H3K27me3 ChIP-seq (EED F97A) | Figure S4 | GSM2475247 | [7] |
| 31 | mESC Replication Timing | Figure 2, 3, 4, 5 | GSE137764 | [8] |
| 32 | H3K27me3 T0 ChOR-Seq | Figure S4 | GSM6203582 | [9] |
| 33 | H3K27me3 T0 ChOR-Seq | Figure S4 | GSM6203583 | [9] |
| 34 | H3K27me3 T180 ChOR-Seq | Figure S4 | GSM6203584 | [9] |
| 35 | H3K27me3 T180 ChOR-Seq | Figure S4 | GSM6203585 | [9] |
| 36 | H3K27me3 T480 ChOR-Seq | Figure S4 | GSM6203586 | [9] |
| 37 | H3K27me3 T480 ChOR-Seq | Figure S4 | GSM6203587 | [9] |
| 38 | H3K27me3 T720 ChOR-Seq | Figure S4 | GSM6203588 | [9] |
| 39 | H3K27me3 T720 ChOR-Seq | Figure S4 | GSM6203589 | [9] |
| 40 | H3K27me3 CUT&Flow | Figure 2, 3, 4, 5, 6, 7 | This study | |

### References


1. Banaszynski, L.A., Wen, D., Dewell, S., Whitcomb, S.J., Lin, M., Diaz, N., Elsasser, S.J., Chapgier, A., Goldberg, A.D., Canaani, E., et al. (2013). Hira-dependent histone H3.3 deposition facilitates PRC2 recruitment at developmental loci in ES cells. Cell *155*, 107-120. 10.1016/j.cell.2013.08.061.
2. Hojfeldt, J.W., Laugesen, A., Willumsen, B.M., Damhofer, H., Hedehus, L., Tvardovskiy, A., Mohammad, F., Jensen, O.N., and Helin, K. (2018). Accurate H3K27 methylation can be established de novo by SUZ12-directed PRC2. Nat Struct Mol Biol *25*, 225-232. 10.1038/s41594-018-0036-6.
3. Perino, M., van Mierlo, G., Karemaker, I.D., van Genesen, S., Vermeulen, M., Marks, H., van Heeringen, S.J., and Veenstra, G.J.C. (2018). MTF2 recruits Polycomb Repressive Complex 2 by helical-shape-selective DNA binding. Nat Genet *50*, 1002-1010. 10.1038/s41588-018-0134-8.
4. Fursova, N.A., Blackledge, N.P., Nakayama, M., Ito, S., Koseki, Y., Farcas, A.M., King, H.W., Koseki, H., and Klose, R.J. (2019). Synergy between Variant PRC1 Complexes Defines Polycomb-Mediated Gene Repression. Mol Cell *74*, 1020-1036 e1028. 10.1016/j.molcel.2019.03.024.
5. de Dieuleveult, M., Yen, K., Hmitou, I., Depaux, A., Boussouar, F., Bou Dargham, D., Jounier, S., Humbertclaude, H., Ribierre, F., Baulard, C., et al. (2016). Genome-wide nucleosome specificity and function of chromatin remodellers in ES cells. Nature *530*, 113-116. 10.1038/nature16505.
6. Goldberg, A.D., Banaszynski, L.A., Noh, K.M., Lewis, P.W., Elsaesser, S.J., Stadler, S., Dewell, S., Law, M., Guo, X., Li, X., et al. (2010). Distinct factors control histone variant H3.3 localization at specific genomic regions. Cell *140*, 678-691. 10.1016/j.cell.2010.01.003.
7. Oksuz, O., Narendra, V., Lee, C.H., Descostes, N., LeRoy, G., Raviram, R., Blumenberg, L., Karch, K., Rocha, P.P., Garcia, B.A., et al. (2018). Capturing the Onset of PRC2-Mediated Repressive Domain Formation. Mol Cell *70*, 1149-1162 e1145. 10.1016/j.molcel.2018.05.023.
8. Zhao, P.A., Sasaki, T., and Gilbert, D.M. (2020). High-resolution Repli-Seq defines the temporal choreography of initiation, elongation and termination of replication in mammalian cells. Genome Biology *21*, 76. 10.1186/s13059-020-01983-8.
9. Flury, V., Reveron-Gomez, N., Alcaraz, N., Stewart-Morgan, K.R., Wenger, A., Klose, R.J., and Groth, A. (2023). Recycling of modified H2A-H2B provides short-term memory of chromatin states. Cell *186*, 1050-1065 e1019. 10.1016/j.cell.2023.01.007.
